# Budding yeast centromeric DNA and A+T rich bacterial DNA can function as centromeres in the fission yeast *Schizosaccharomyces pombe*

**DOI:** 10.1101/513150

**Authors:** Anne C Barbosa, Zhengyao Xu, Kazhal Karari, Silke Hauf, William RA Brown

**Affiliations:** School of Life Sciences, University of Nottingham, Queen’s Medical Centre, NG7 2UH, UK; Department of Biological Sciences, Biocomplexity Institute, 1015 Life Science Circle, Blacksburg, VA 24061, USA

**Author notes:** Contributed equally to the work described and wish to be acknowledged as joint first authors (Anne C Barbosa, Zhengyao Xu). Current addresses; ACB; Laboratory of Human Genetics, Department of Genetics and Evolutionary Biology, Institute of Biosciences of the University of São Paulo - USP Rua do Matão, n°. 277, Cep: 05508-090, Cidade Universitária - São Paulo – SP, Brazil. KK; Department of Biology, College of Sciences, Raparin University, KRG-IRAQ.

## Abstract

Eukaryotic centromeric DNA is famously variable in evolution but currently, this cannot be reconciled with the conservation of eukaryotic centromere function. It seems likely that centromeric DNA from different organisms contains conserved functionally important features but the identity of these features is unknown. The point centromeres of the budding yeast *Saccharomyces cerevisiae* and the regional centromeres of the fission yeast *Schizosaccharomyces pombe* are separated by 350 million years of evolution and are canonical examples of the paradoxical relationship^1^ between centromeric DNA sequence and function. We have established a centromere-replacement strategy in *Schizosaccharomyces pombe* in order to resolve this paradox experimentally. Centromere-replacement shows that an A+T rich bacterial DNA sequence has weak centromere function and that elements of the *Saccharomyces cerevisiae* centromere embedded in short sequences from the non-centromeric *S. pombe wee1* gene function almost as well as native *S. pombe* centromeric DNA. These observations demonstrate that determinants of centromere function are held in common by the budding and fission yeasts and that A+T rich DNA is both necessary and sufficient for function in *S. pombe*. Given the evolutionary distance between these yeasts, it is likely that A+T rich DNA has centromere function in a wide variety of eukaryotes. Centromere-replacement uses unidirectional serine recombinases that work well in many organisms^2 3^ and our experimental strategy should allow this idea to be tested in other eukaryotes.

---

The fission yeast *Schizosaccharomyces pombe* is ideal for studying centromeric DNA because it possesses a sophisticated set of genetical tools and centromeres that are similar to the regional centromeres of many eukaryotes^4 5^. However, these potential advantages have never been fully exploited because there has been no easy way of studying centromeric DNA in a chromosomal context. Instead the function of centromeric DNA in *S. pombe* has been studied using plasmid-based approaches^6,7^, which suffer from variable copy numbers, uncertain structures and from difficulties of measuring centromere function because plasmids show an elevated rate of loss compared to natural linear chromosomes^8^. These problems would all be avoided if native centromere DNA could be replaced with candidate DNA. However, there have been two obstacles to implementing such a simple approach. The first is that, until recently, there has been no easy way of serially integrating and deleting DNA. The second is that the centromeres of the commonly-used laboratory strain of *S. pombe* are flanked by palindromically organized heterochromatin that makes sequence manipulations difficult. The first limitation was overcome with the development of a set of functionally optimized unidirectional serine recombinases^2 3^. That the second limitation may also no longer apply was suggested by the discovery of a natural isolate of *S. pombe* (CBS2777)^9^ that has two centromeres, those of chromosomes 2 and 4^10^, which lack flanking heterochromatin. Live cell imaging of CBS2777 with a tdTomato-labelled chromosome 2 and GFP-labelled centromeric histone CENP-A^Cnp1^ showed that most mitoses were accurate although anaphase was delayed by about 5 minutes relative to the laboratory strain (Supplementary figure 6). Despite the importance of heterochromatin for binding cohesin and preventing merotelic attachment^11, 12^, there was no detectable excess of lagging chromosomes. We concluded that the heterochromatin-deficient centromere on chromosome 2 of CBS 2777 is largely functional and have therefore used it for centromere-replacement experiments.

We established a two-step approach to centromere-replacement (Figure 1A in outline, and Supplementary data in detail). ϕC31 integrase was used to place a candidate sequence or an empty vector adjacent to the native centromere of chromosome 2 of CBS2777, and Bxb1 integrase was subsequently used to delete the native centromere. When centromere central core DNA was used as a candidate sequence, cells that had deleted the native centromere were recovered according to the length of the candidate DNA. Full-length (9.5kb) central core DNA led to recovery with approximately 100% efficiency (Figure 1B) whereas central core sequences less than 3.58kb showed approximately 0.1% recovery, the same as an empty integration vector. This level of recovery was consistent with the rates of neocentromere formation^5^ and, as we show below, such deleted cells do contain neo-centromeres. The three orders of magnitude dynamic range in the efficiency of recovery of cells that had successfully deleted the native centromere meant that the recovery of such cells could be used to compare the abilities of different candidate sequences to support centromere function.

**Figure 1.**
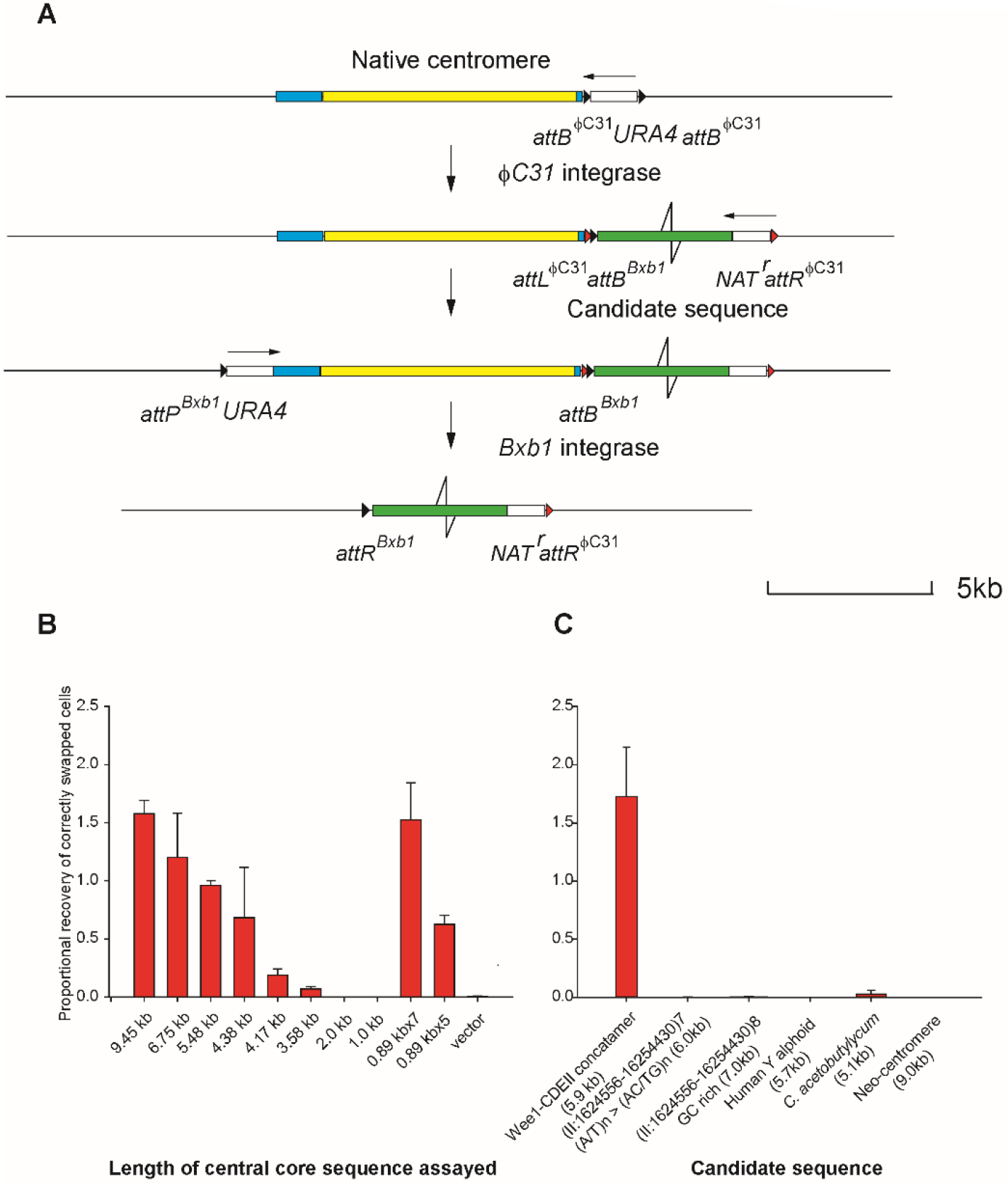
Centromere-replacement in *S. pombe* CBS 2777 as an assay for the centromeric activity of centromeric DNA. **A.** Sequence manipulations involved in centromere-replacement on chromosome 2 in CBS 2777. For a summary see the main text and for details see the Supplementary data. **B.** Recovery of cells that had deleted the native centromeric DNA following transfection with a Rep81-Bxb1 integrase expression plasmid as a proportion of cells following transformation with an empty Rep81 expression vector for strains in which the different centromere sequences indicated in B were placed adjacent to the native centromere. The data are summaries of the mean and standard deviations of the results detailed in Supplementary data table 4. The 0.89kb sequence was concatamerized into pentamers or heptamers of 4.4kb and 6.2kb The assay is described in detail in the Supplementary data which also discusses the observation that the maximum proportional recovery is greater than 1. **C.** Assay of different candidate sequences by centromere-replacement. The data are summaries of the mean and standard deviations of the results detailed in Supplementary data table 4. These data represent the recovery of cells that had deleted the native centromeric DNA upon transformation with the Rep81-Bxb1 expression vector expressed as a proportion of the recovery of cells upon transformation with an empty Rep81 expression vector.

Concatamers of five or seven copies of an 889 bp sub-section of the central core were as functional as intact central core DNA of similar length, suggesting that the functionality of longer pieces is not due to complementary unrelated activities, but to the repetition of features that are present in shorter segments (Figure 1B, Supplementary Table 4). The dependence of centromere formation upon the length of central core DNA is consistent with earlier results^7^ using plasmid-based assays.

The experimental centromere sequences were derived from the laboratory strain of *S. pombe* and therefore differed from the CBS2777 strain centromere 2 sequences by indels and SNPs^9 13^. They had also been marked by the introduction of 4bp deletions of random sequences approximately every 1kb by mutagenic PCR. Overall, the experimental sequences were 94.9% identical to the centromeric DNA of chromosome 2 of CBS 2777 and 98.9% identical to the centromeric central core of chromosome II of the laboratory strain. These differences allowed discrimination between the experimental sequences and the native CBS2777 centromeric DNA and enabled ChIP-seq to measure binding of centromere proteins (Figure 2A and B for the 9.5kb sequence and Supplementary figure 8 for the 6.75kb sequence and for the concatamers of the 889bp sequence). This showed that the experimental sequences bound CENP-A^Cnp1^ prior to replacement and that this pattern was retained after deletion of the native centromere. The ChIP-seq data were normalized by comparison to the total number of chromosome 1 and 3 reads and the normalized read depth showed an overall increase in the extent of the CENP-A^Cnp1^ domain in the presence of functional test sequences. We also carried out ChIP-seq for CENP-C^Cnp3^ prior to replacement of the 9.5kb test sequence. The pattern (Figure 2C) of CENP-C^Cnp3^ binding matched that for CENP-A^Cnp1^, suggesting that the entire kinetochore was assembled, consistent with the viability of centromere-replaced cells.

**Figure 2.**
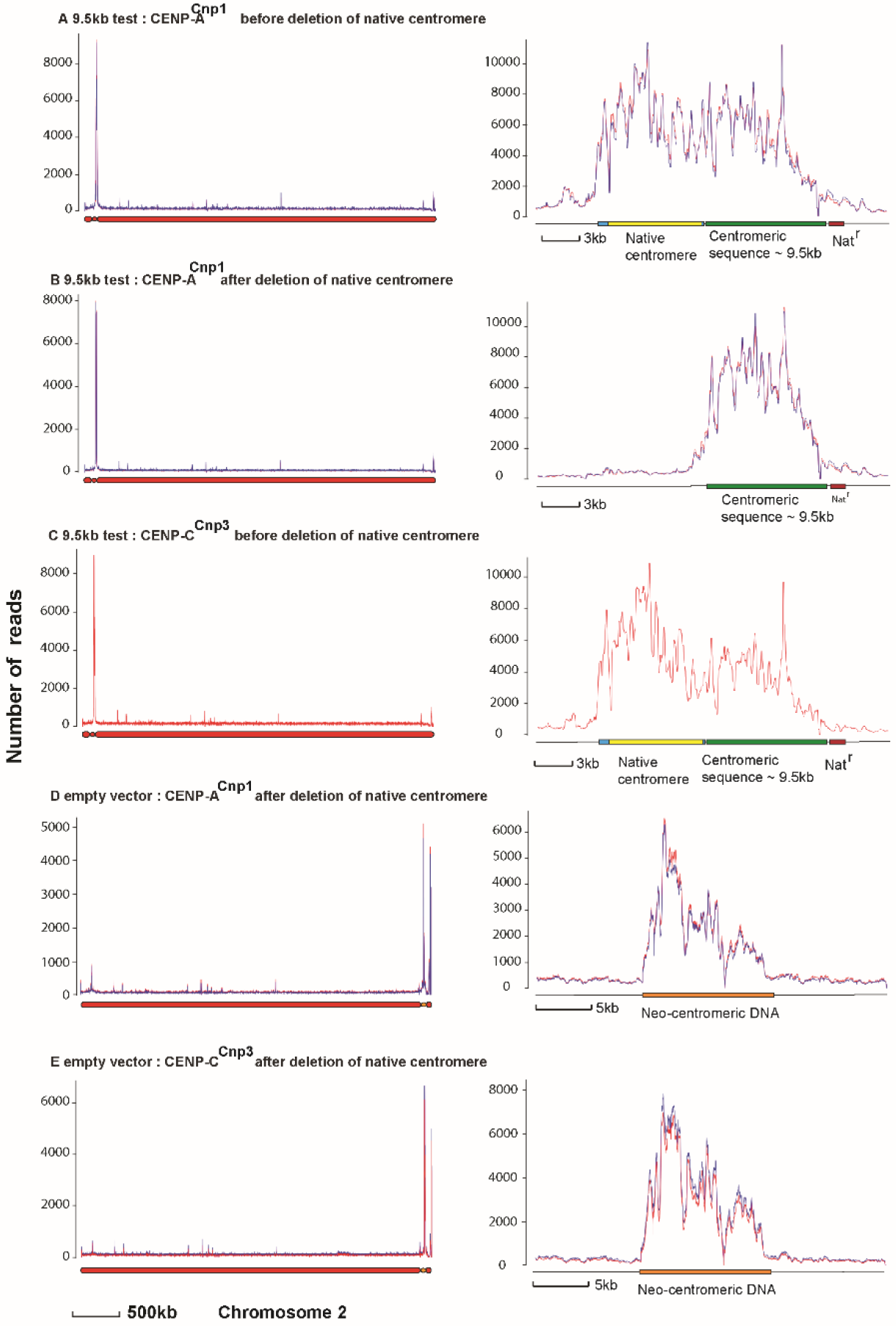
CENP-A^Cnp1^ and CENP-C^Cnp3^ binding to centromere and neo-centromere sequences before and after centromere-replacement in *S. pombe* CBS 2777. **A, B.** Binding of CENP-A^Cnp1^ to an intact 9.5kb centromere central core placed adjacent to the native centromere of CBS 2777 chromosome 2 before (A) and after (B) replacement. In this, and in all subsequent parts of the figure, the left panel shows binding to the whole chromosome; the right panel to the centromere region. **C.** Binding of CENP-C^Cnp3^ to an intact 9.5kb centromere central core placed adjacent to the native centromere of CBS 2777 chromosome 2 prior to replacement. **D, E.** Binding of CENP-A^Cnp1^ and CENP-C^Cnp3^ to the neo-centromere region of CBS 2777 chromosome 2 after deletion of the native centromere in a strain with an empty vector adjacent to the native centromere

In clones where empty vector sequence had been placed adjacent to the native centromere prior to deletion, all of the CENP-A^Cnp1^ was located on distal sub-telomeric DNA (Figure 3D) and mainly on a 13kb stretch of DNA approximately 70kb from the telomere. This stretch of DNA also bound CENP-C^Cnp3^ (Figure 2E) in a similar pattern consistent with neo-centromere formation^5^. In total, we analysed six centromere-deleted clones derived from the empty vector or from experimental sequences less than 3.58kb that showed no detectable activity (2.0 kb and 1kb candidates), and all showed CENP-A^Cnp1^ binding to the same distal sub-telomeric stretch of DNA.

**Figure 3.**
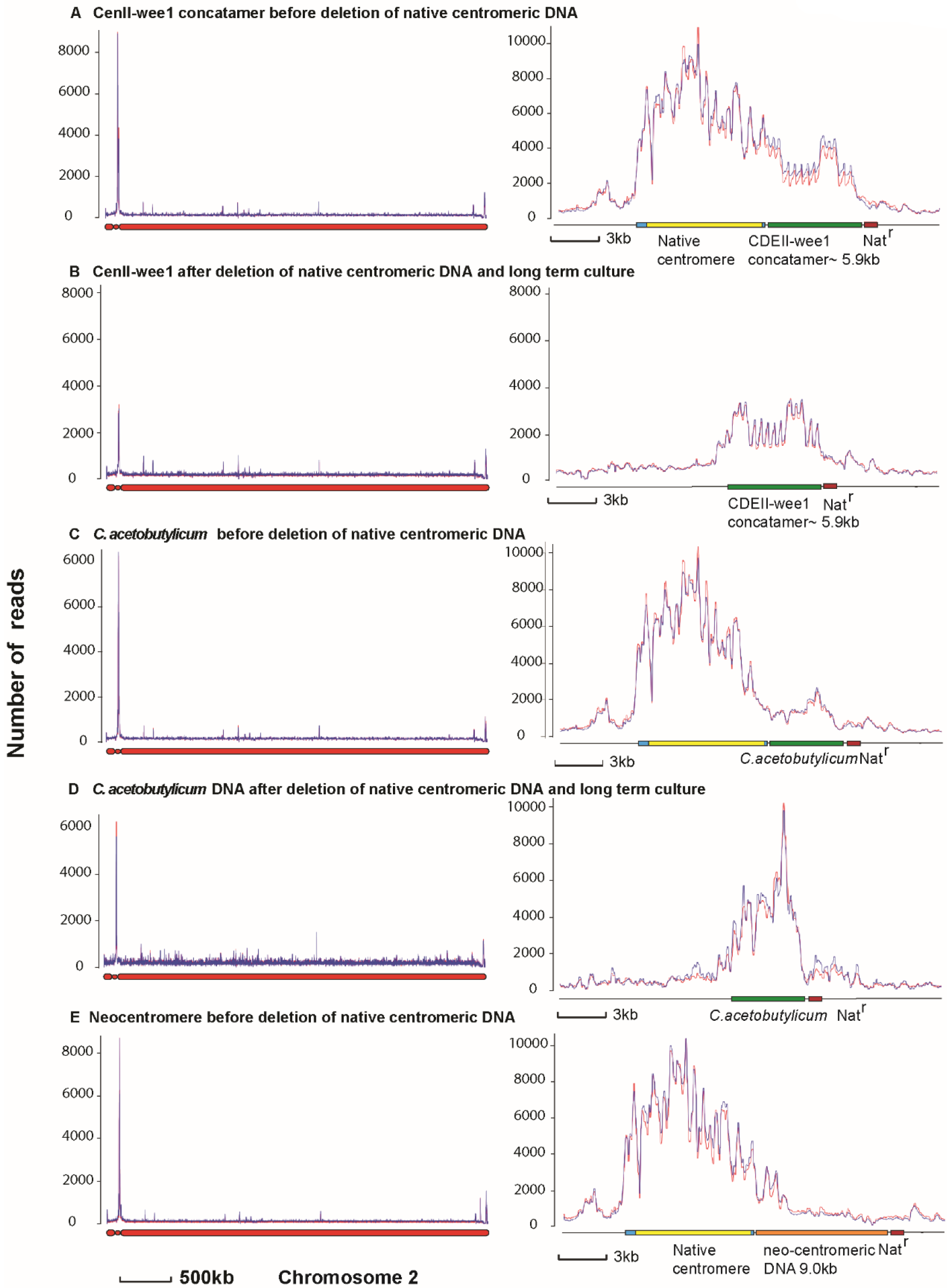
CENP-A^Cnp1^ binding to candidate sequences before and after centromere-replacement in *S. pombe* CBS 2777. **A, B.** Binding of CENP-A^Cnp1^ to a CDEII-wee1 concatatemer placed adjacent to the native centromere of CBS 2777 chromosome 2 before and after replacement. In this, and in all subsequent parts of the figure, the left panel shows binding to the whole chromosome; the right panel to the centromere region. **C, D.** Binding of CENP-A^Cnp1^ to the 5.1kb stretch of *C. acetobutylicum* DNA when placed adjacent to the native centromere of CBS 2777 chromosome 2 before and after replacement. **E.** Binding of CENP-A^Cnp1^ to a 9kb stretch of CBS 2777 chromosome neo-centromeric DNA when placed adjacent to the native centromere of CBS 2777 chromosome 2.

The two shortest of the central core sequences to show clone recovery above background in the centromere-replacement assay were 4.17kb and 3.58kb. In order to know whether such short centromeres were capable of supporting stable centromere function we pooled 40 clones derived from these sequences into two groups of twenty and analysed these two pools by ChIP-seq for CENP-A^Cnp1^. The replacement clones containing the 4.17kb sequence showed CENP-A^Cnp1^ binding exclusively to the candidate sequence (supplementary figure 8C) and we concluded that centromeric DNA of this length was functional. To test whether these centromeres were stable, we cultured 20 sub-clones in an approach analogous to that used in mutation accumulation experiments and designed to avoid the effects of selection^14 15^. There was no major difference between CENP-A^Cnp1^ binding before and after culture and, in particular, no detectable neo-centromere formation (supplementary figure 8D). Given a colony size of 5 x10^6^ at each round of streaking, these results indicate that the 4.17kb centromere is stable with a rate of conversion from the centromere to the neo-centromere of less than 1.5% per cell division (with 95% confidence). The replacement clones containing the 3.58kb centromere sequence grew slowly and we were initially unable to derive ChIP-seq data. After several months storage with two rounds of restreaking to ensure viability, the clones had recovered growth and showed an exclusively neo-centromeric pattern of CENP-A^Cnp1^ binding suggesting that centromeric sequences of 3.58kb are unable to support the stable presence of a centromere.

To analyze whether CENP-A^Cnp1^ binding to the modified centromeric test sequences was contingent upon the initially juxta-centromeric location, we placed the 9.5kb stretch of centromeric central core at either the *ade6* locus on chromosome 3 (214.3 kb from the centromere) or at the chromosome 2 neo-centromere locus described above. ChIP-seq for CENP-A^Cnp1^ showed that neither ectopically placed sequence bound detectable CENP-A^Cnp1^ (supplementary figure 9), and thus CENP-A^Cnp1^ binding to a candidate sequence is contingent upon the juxta-centromeric location. Seeding by the natural centromere thus explains the high recovery of replacement clones in our assay compared to the extremely low frequency of neo-centromere formation.

Our assay is ideal for assessing which features of centromeric DNA sequences are important for functionality because it is sensitive and because it tests function at a centromere location. Centromere sequences are characteristically A+T-rich with respect to the rest of the genome. Native *S. pombe* centromere central core sequences together with their imr repeats contain between 70.6 and 71.9% A+T compared with a genome average of 63.9% A+T, and human Y alphoid DNA contains 63% A+T compared with a genome average of 53.9% A+T. To test for the importance of A+T content, we mutated the 889 bp sequence that we had shown to be active as a tandem array of five or seven copies (Figure 1C). We either disrupted the A/T runs in the sequence with alternate Cs or Gs or introduced Cs or Gs outside the A/T runs (supplementary data section 2.2). Both approaches reduced the A+T content from approximately 70% to 61 % and destroyed the centromere activity of the sequence as well as its ability to bind CENP-A^Cnp1^ (Figure 1C, supplementary figure 10). Hence, A/T runs are insufficient and an overall high A+T content is required. Consistently, a 5.7kb unit repeat of human Y chromosome alphoid DNA showed less than 1% the activity of an equivalent length of *S. pombe* centromere DNA and no binding of CENP-A^Cnp1^ (Figure 1C, supplementary figure 10).

In order to determine whether A+T content was sufficient for centromere function, we tested a 5.1kb stretch of bacterial *Clostridium acetobutylicum* DNA, which contained 70% A+T (67-73% per kb). The sequence showed weak but detectable activity (Figure 1C) and bound CENP-A^Cnp1^ both before and after the swap (Figure 3C and D). There were no other sites of CENP-A^Cnp1^ binding on chromosome 2 and we therefore concluded that the bacterial DNA functions as a centromere in *S. pombe.* However, even after replacement the binding was lower than observed with *S. pombe* centromeric test sequences of similar length (Figure 4). That this sequence showed lower levels of activity than native centromeric DNA of similar length and base composition established that aspects of centromeric DNA other than A+T content are functionally important.

**Figure 4.**
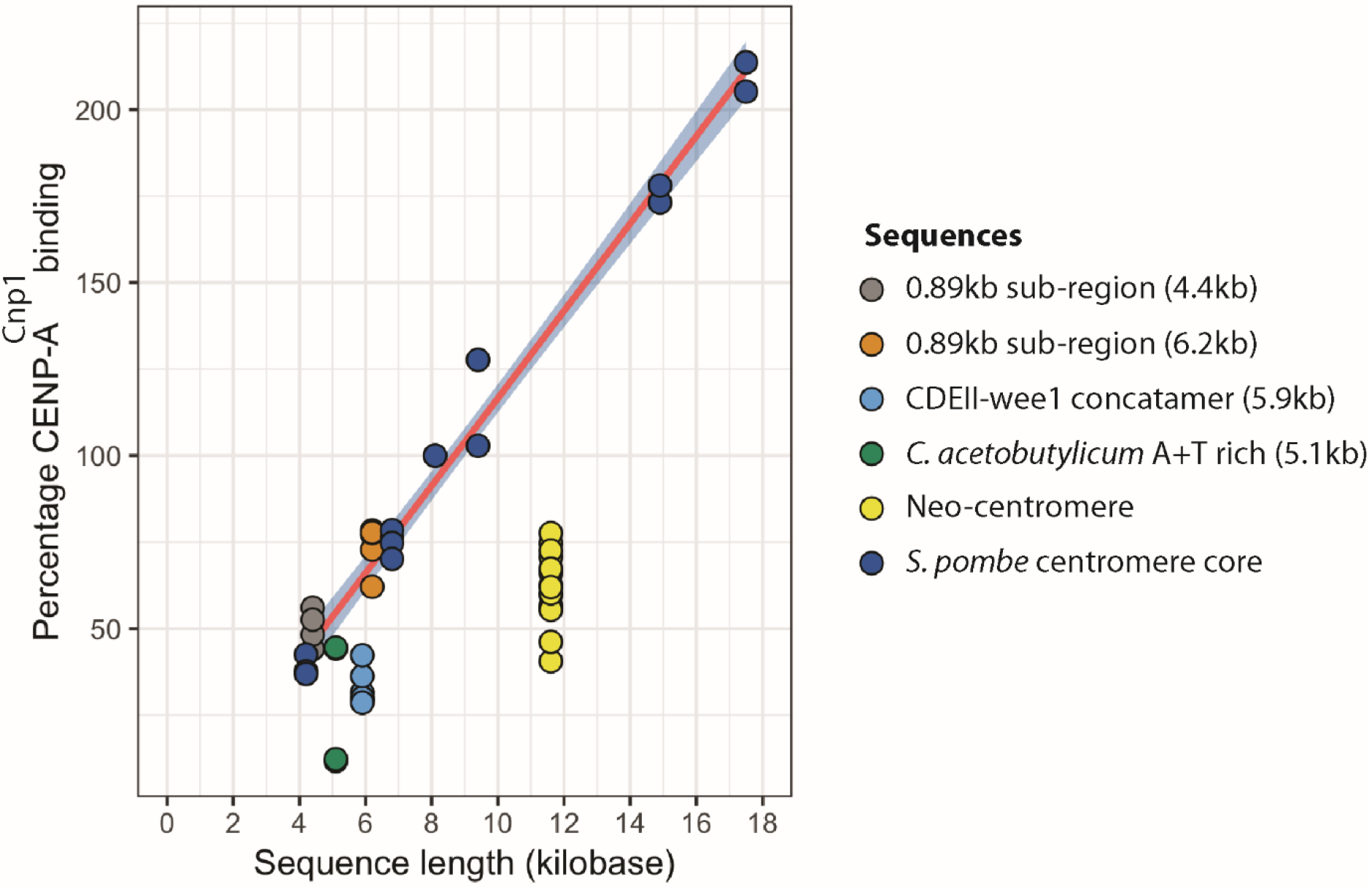
CENP-A^Cnp1^ binding to candidate sequences as a function of sequence length Binding of CENP-A^Cnp1^ to different sequences that functioned as centromeres as a function of their length. The pink line is the regression line of the binding of CENP-A^Cnp1^ to sequences derived from the central core of *S. pombe* centromeric DNA. The grey shading indicates the 95% confidence limits of the regression line. The two data points corresponding to the *C.acetobutylicum* DNA represent the binding before (lower point) and after replacement. For the remaining sequences, no obvious differences were observed between the data before and after replacement. The data are normalized to the intact central core DNA that is 9.5kb.

The neo-centromere at the sub-telomere of CBS2777 chromosome 2 has an A+T content of 68%. However, when analysed in the centromere-replacement assay, the neo-centromeric DNA showed less than 1% of the activity of an equivalent length of *S. pombe* centromeric DNA (Figure 1C). Before replacement CENP-A^Cnp1^ binding extended only about 1kb into the neo-centromeric DNA. (Figure 3E) rather than throughout the sequence as seen at the neo-centromere. This assay also demonstrates that the sequence of a neo-centromere and the ability to bind CENP-A^Cnp1^ are insufficient to support centromere function. Neo-centromeres are typically found adjacent to heterochromatin, and our findings are consistent with the possibility that the presence of heterochromatin in *cis* is important in acquiring neo-centromere function. Alternatively, factors only expressed in cells with neo-centromeres may be acting in *trans* to allow neo-centromere sequences to function as centromeres^5^.

We have established a powerful assay to test for features of centromeric DNA required at an endogenous centromere locus. Since bacterial DNA established a centromere in our assay, albeit with low efficiency, we wanted to test DNA from other eukaryotes. *S. pombe* and the budding yeast *Saccharomyces cerevisiae* diverged approx. 350 million years ago and contain fundamentally different centromeres^16^. *S. cerevisiae* contains ‘point’ centromeres defined by 125 bp sequence (consisting of CDEI, II and III)^17^ and a single CENP-A^CSE4^-containing nucleosome^18^ while *S. pombe* contains regional centromeres with several CENP-A^Cnp1^-containing nucleosomes. To test *S. cerevisiae* centromere DNA functions in *S. pombe*, we embedded 20-25 bp stretches of *S.cerevisiae* CDEII elements derived from different chromosomes into short blocks of the *S. pombe wee1* gene, which was arbitrarily chosen as a non-centromeric sequence. Together, these sequences formed a concatamer of 5.9 kb with 74% A+T content (supplementary figure 7). Strikingly, centromere-replacement demonstrated that the CDEII-concatamer showed activity that was similar to native *S. pombe* centromeric DNA (Figure 1B, C). CENP-A^Cnp1^ bound to the CDEII-concatamer sequence both before and after replacement (Figure 3A and B) albeit at levels that were lower than for *S. pombe* central core centromeric DNA (Figure 4). Hence *S. cerevisiae* CDEII can support centromere function in *S. pombe* when assembled into a repetitive array of adequate length. This indicates that despite their divergence, budding and fission yeasts share a mechanism to define centromeres.

Overall, our centromere-replacement assay has demonstrated the importance of A+T content for the function of centromeric DNA while highlighting the significance of other characteristics for the native centromere itself. These facts can be reconciled by a model, similar to those used to account for patterns of codon bias^19^, in which the sequences of native centromeres are maintained by a balance between selection in favour of A or T residues and mutation in the direction of the bulk genome composition. Strikingly, A+T-rich DNA from bacteria was able to establish a centromere in *S. pombe*, whereas an *S. pombe* neo-centromere was unable to replace the centromere at the endogenous locus. The sequence features that allow or inhibit neo-centromere formation beyond a general A+T richness have not been identified, but our assay will allow testing the possibilities. Our centromere-replacement strategy combines the activities of two well characterized and broadly efficient serine integrases^2 3^, and should enable the testing of the centromere functionality of a range of DNA sequences not only in *S. pombe* but also in a variety of other eukaryotic species. Thus it will allow a resolution of the so-called “centromere-paradox”^1^.

## Supporting information

Entire_supplementary_data

## Acknowledgements

We thank Takeshi Sakuno and Yoshinori Watanabe for their gift of the tetO array/td-Tomato system. We also thank Sheila McCormick, Jonas Warringer and Marshall Stark for comments. The work in Nottingham was supported by BBSRC BB/K003356/1. ACB was funded by the Brazilian ‘Nottingham-Birmingham’ PhD scheme organised by CAPES, Brazil. KK was funded by the Kurdistan Regional Government, Human Capacity Development Program. SH was supported by the National Institute of General Medical Sciences of the National Institutes of Health (R35GM119723).

