## Supplementary material for "Budding yeast centromeric DNA and A+T rich bacterial DNA can function as centromeres in the fission yeast *Schizosaccharomyces pombe*": Entire_supplementary_data

**Barbosa et al., Supplementary data**

### 1 Chromosome engineering

#### 1.1 Engineering a chromosome to enable assay of the centromere activity of a candidate sequence.

Our approach to assaying the ability of a particular sequence to function as a centromere consisted of two main stages. The first involved the construction of a chromosome that contained the candidate centromeric sequence placed adjacent to the pre-existing centromere which was in turn flanked by attachment sites for the Bxb1 integrase (supplementary data figure 1). This was achieved through the following three steps:

1. A Ura4 gene flanked by attachment B sites for the ϕC31 integrase was integrated between residues 131490 and 131491 adjacent to breakpoint 4 sequence on the right-hand side of the centromere of chromosome 2 of CBS 2777 Ura4$^{\Delta}$ Leu1$^{\Delta}$ kanMX6-Pcnp1-mEGFP-cnp1(Nott363) to create strain Nott 373 (Supplementary data, table 2). The targeting construct was assembled using primers 420-423 (Supplementary data, table 3). Targeted integration was assayed by PCR and mapped to chromosome 2 by pulsed-field gel electrophoresis and filter hybridization.
2. Candidate sequences (see below) were cloned into the BamHI site of the vector pFA6a-natMX6 REV *attP*$^{\phi C31}$ *attP* $^{\phi C31}$ attB$^{Bxb1}$ (supplementary data figure 2) and then integrated into the attB sites of breakpoint 4 construct in strain Nott 373 using ϕC31integrase expressed from a pREP1 construct encoding a codon-optimized form of the gene tagged with a nuclear localization sequence. Integration of the candidate sequence cloned in pFA6a-natMX6 REV *attP*$^{\phi C31}$ *attP* $^{\phi C31}$ attB$^{Bxb1}$ was accompanied by loss of the Ura4 marker and acquisition of resistance to the antibiotic nourseothricin which was used as the basis of an initial screen for site-specific integration. The empty pFA6a-natMX6 REV *attP*$^{\phi C31}$ *attP* $^{\phi C31}$ attB$^{Bxb1}$ vector was integrated as a control. Integration was then checked by PCR at both ends of the integrated DNA and the integrity of the integrated sequences was checked by long-range PCR and, in the early experiments, by agarose gel electrophoresis, filter transfer and hybridization. In the initial experiments aimed at establishing the system, site-specific recombination was also confirmed as reciprocal and conservative by recovering and sequencing the ϕC31integrase *attL* and *attR* sites (Anne Barbosa, PhD thesis University of Nottingham, 2017).
3. The left-hand side (breakpoint 3) of the centromere of each strain containing a candidate sequence was then targeted between positions 123238 and 123281 with a Ura4 gene flanked by attachment P site for the Bxb1 integrase. This construct was engineered using primers 880, 982, 882 and 883. The centromeres of chromosomes 2 and 4 are homologous and so it was necessary to screen for targeting and then to determine the chromosomal location of the targeted construct. The targeting screen was carried out using primers pairs 893/884 and 892/885 (Supplementary data, table 3). The chromosomal mapping was carried out by long-range PCR using primer 893 and either 1030 which was complementary to the ϕC31integrase *attL* site on chromosome 2 or primer 106 which was complementary to the un-engineered chromosome 4.

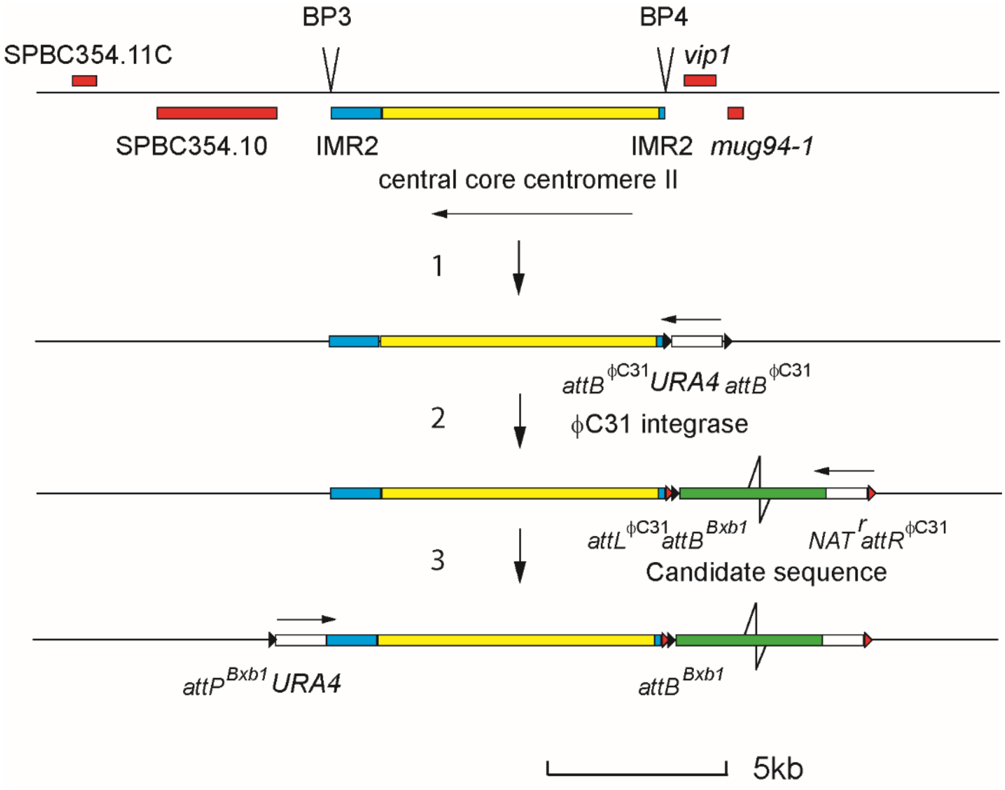


**Supplementary data figure 1**. Sequence of steps used to engineer a chromosome containing a candidate sequence adjacent to the native centromere

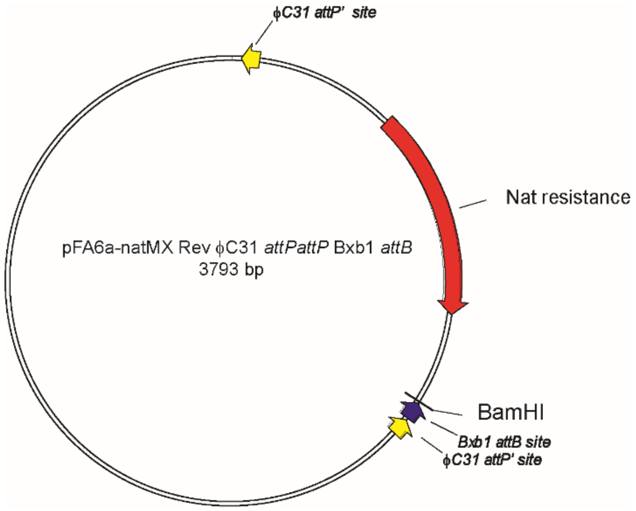


**Supplementary data figure 2**. pFA6a-natMX6 REV *attP*$^{\phi C31}$ *attP* $^{\phi C31}$ attB$^{Bxb1}$ vector used to integrate sequences into chromosomes using site specific recombination with ϕC31integrase

#### 1.2 Assaying the ability of a candidate sequence to support centromere formation using site-specific recombination.

The assay of the ability of a sequence to function as a centromere consisted of the deletion of the pre-existing centromeric DNA using the Bxb1 integrase (supplementary data figure 3) and measurement of the efficiency with which cells that had deleted the native centromere were recovered.

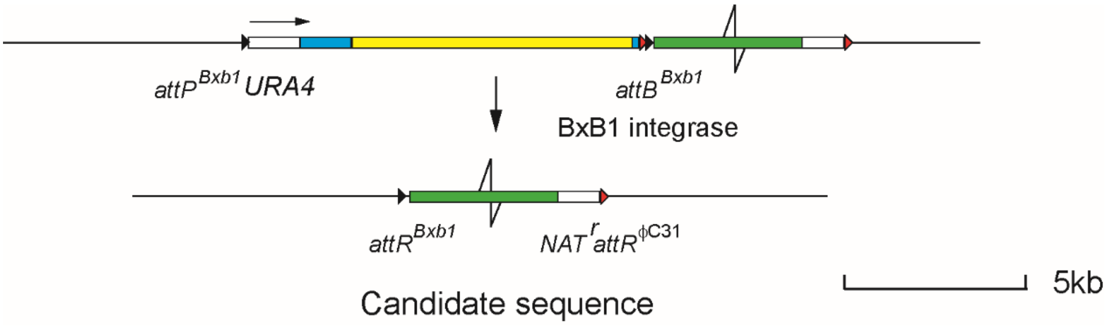


**Supplementary data figure 3**. Centromere replacement using Bxb1 integrase

In detail the assay procedure consisted of the following steps;

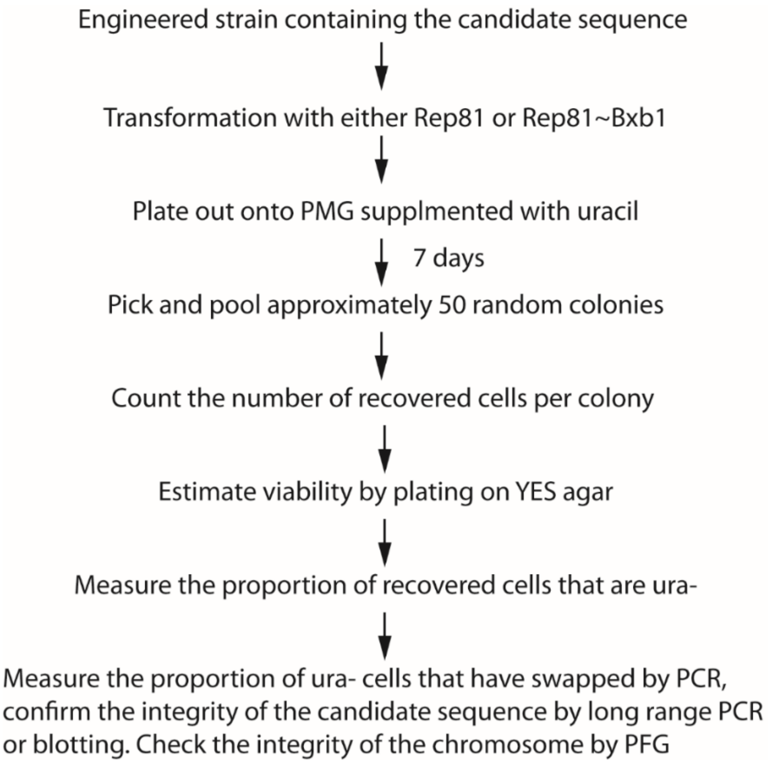


1. The first step was to delete the native centromere using Bxb1 integrase. This involved transformation either with a Bxb1 integrase expression construct; pRep81Bxb1 integrase or, as a control, an empty vector pRep81. The details of the transformation procedure are given below. After transformation, the cells were allowed to recover while selecting for the presence

of the expression plasmid and maintaining uracil in the medium to allow cells that have lost the native centromere and flanking Ura4 marker to proliferate.

2. After seven days at 32°C the transformation plates were removed from the incubator. The yield of colonies was variable. Typically between 25 and 300 colonies were recovered in a single transformation but no systematic differences in the numbers of transformed colonies were observed following transformation of cells containing engineered chromosome 2 with an entire central core as a candidate sequence and cells containing engineered chromosome 2 with an empty vector as a candidate sequence. However, the average size of the colonies recovered upon transformation of the cells containing an empty vector were smaller than upon transformation of the cells containing an entire central core sequence (supplementary data; figure 4 and table 4).

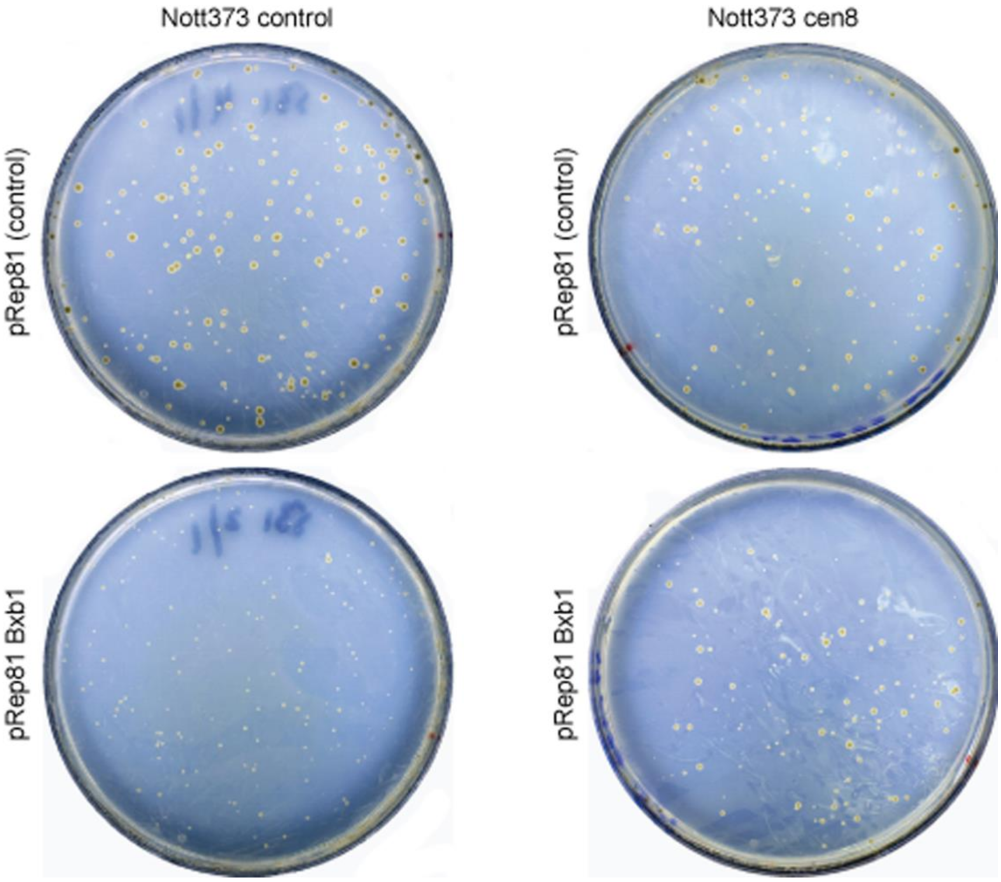


**Supplementary data figure 4**. Differences in colony size between cells containing an intact central core as a candidate sequence and cells containing an empty vector upon transformation with pREP81 Bxb1 expression construct. Nott 373 control contained the empty pFA6a-natMX6 REV $attP^{\phi C31}$ $attP^{\phi C31}$ attB$^{Bxb1}$ vector while Nott373 cen8 contained an intact central core as a candidate sequence. Cells of either of these two strains were transformed with either the empty pREP81 expression plasmid or the pREP81 Bxb1 expression vector and recovered on PMG supplemented with uracil. Colonies that on average are smaller are specifically recovered upon transformation of Nott373 control with the pREP81 Bxb1 expression vector (lower left plate).

3. The difference between the sizes of the colonies recovered following transformation of a strain containing the empty pFA6a-natMX6 REV $attP^{\phi C31}$ $attP^{\phi C31}$ attB$^{Bxb1}$ (supplementary data; figure 4) with the Rep81 and the Rep81Bxb1 plasmids could be explained by the fact that the integrase was excising the native centromere of chromosome 2 at a rate of less than once per cell cycle and that the cells lacking the centromere were either dying or forming a neo-

centromere at a low frequency. In the cells containing an intact central core as a candidate sequence, the loss of the native centromere was compensated by the candidate sequence. In order to test this idea and to assay the functionality of a sequence we picked and pooled approximately 50 colonies from the cells transformed with either the empty pREP81 plasmid or the Rep81Bxb1 expression plasmid, counted the number of cells in the pool to yield an estimate of the average number of cells in each colony. We then estimated the viability of these cells by plating out a known number (-1000 in duplicate or triplicate) on to YES agar and counting the number of recovered colonies. Of these, a fraction dependent upon the nature of the candidate sequence should have successfully deleted the native centromere and be Ura-. We estimated this fraction by patching onto YES and PMG + Leu agar and recording the proportion that failed to grow in the absence of uracil. Finally, we measured the proportion of these cells that had accurately deleted the native centromere by PCR (supplementary data; table 4)

4. In at least ten ura- clones, we confirmed the integrity of the candidate sequence and the accuracy of the replacement by long-range PCR or restriction enzyme digestion, gel electrophoresis, filter transfer and hybridization (supplementary data figure 5). Similarly, we checked at least ten clones of each type for their chromosomal integrity by pulsed-field gel electrophoresis. The figures used to produce the data used in the bar charts shown in figures 3 and 5 of the main text are given separately in the supplementary data table 4.

The positive control in the assay consisted of an intact 9.45kb central core of centromeric DNA. Using this sequence however we note a discrepancy between what we had expected and what we observed; thus as indicated we recovered in figure 1 of the main text about 60% more deleted cells when we transformed with the Rep81~Bxb1 integrase expression plasmid than the number of centromere intact cells that were recovered after transformation of the same strain with Rep81 alone. The most obvious explanation for the discrepancy is that the cells with the extra copy of the central core sequences grow more slowly than those with a single copy. However, we excluded this explanation by measuring colony sizes of deleted and intact cells after plating onto agar plates. We, therefore, suggest that the difference in colony sizes reflect a difference in the rates of recovery between the cells with just a single copy of the candidate sequence and those with two following transformation. Despite this discrepancy comparison between the recoveries of centromere deleted cells with the different sequences allows a rank-ordered comparison of the different sequences although the comparison is unlikely to be proportional to the ability of the different sequences to support centromere formation after centromere deletion at the single-celled level given that we are uncertain about the rates of proliferation of the centromere deleted and centromere intact cells with different amounts of candidate sequence.

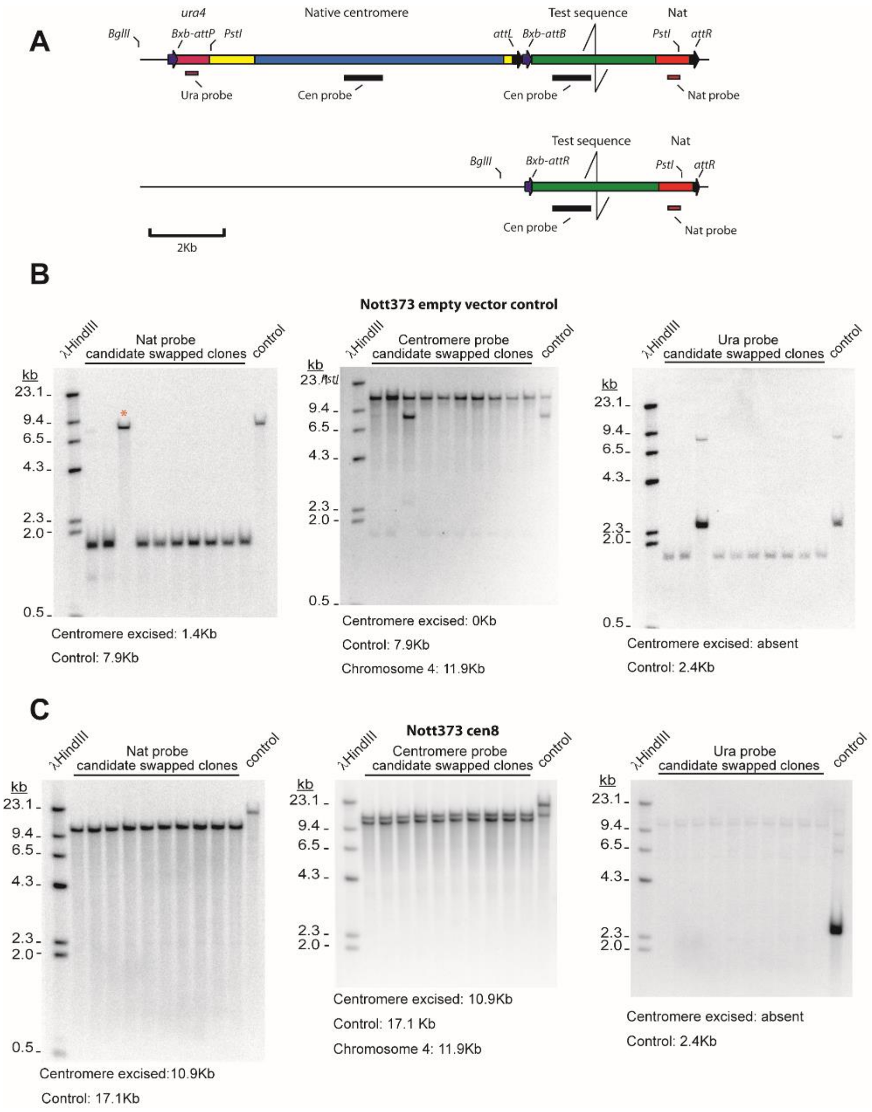


**Supplementary data figure 5.** Checking the sequence organization of the centromeric region of the empty vector and 9.5kb central core (Nott 373 cen8) containing clones before and after deleting the native centromere. **A**; Sequence organization of the DNA with relevant restriction enzyme sites. **B** and **C**; Filter hybridization analysis of the clones before (labelled control) and after deletion. Sizes of predicted restriction fragments are indicated. The third empty vector candidate deletion clone marked with an asterisk is an example of a re-arranged clone where the replacement had failed.

#### 1.3 Transformation of cells for assay of centromere activity.

The standard transformation procedure was designed to be used for assaying the ability of a candidate sequence to support centromere formation. Cells engineered as above were grown up to an OD of approximately 1 at 600nm in 150 ml of a transformation medium consisting of 5mM Potassium hydrogen phthalate, 5mM $Na_2HPO_4$, 8.5mM Glutamic Acid, 7.3mM glucose, 2.4mM $KH_2PO_4$, 1.35mM $MgSO_4.7H_2O$, 1.14mMNaCl, 4.5mM $CaCl_2.2H_2O$, 25mM $(NH_4)_2SO_4$, 2.44mM $CH_3CO_2K$ supplemented with 1ml of vitamin stock and 0.1ml of mineral stock http://www-bcf.usc.edu/-forsburg/media.html. Leucine and uracil were added to 0.25g/l. Cells were washed with sterile distilled water and once with 0.1M $CH_3CO_2Li$, 10mM Tris, HCl, 1mM EDTA, pH 8.0 (TE). The cells were then re-suspended in 1ml of 0.1M $CH_3CO_2Li$, TE and aliquoted into each of four 1.5ml plastic tubes and then pelleted in bench top centrifuge at 1500g for four minutes and the supernatant was removed. The cell pellet at this stage was typically between 100µl and 150µl in volume. 1.62 mg of DNA in 180 µl of 0.1M $CH_3CO_2Li$, TE was added to each tube to yield a suspension that was between 280 µl and 330 µl in volume. The Rep81 Bxb1 integrase expression plasmid was added to two of the tubes and the empty vector to the other two tubes. The tubes were vortexed to re-suspend the cells in the DNA and then 2.36 volumes of 40% PEG4000 in 0.1M $CH_3CO_2Li$, TE was added to each, the resulting suspensions were again mixed by vortexing and then incubated at 32°C for one hour while being slowly inverted using a rotor. 0.39 volumes (with respect to the mixture of cells and DNA) of dimethyl sulphoxide was then added, the mixtures were vortexed and the tubes placed in a water bath at 40°C for six minutes. The tubes were then filled with YES medium (yeast extract with supplements) http://www-bcf.usc.edu/-forsburg/media.html and the cells concentrated by bench top centrifugation. The cells were then re-suspended in YES and the contents of the two tubes containing the cells transformed with one or other of two plasmids were dispersed into 50ml of YES and allowed to recover for two hours at 32°C with shaking. The cells were then pelleted, rinsed in 50ml of distilled water and plated onto three 9cm plates of pombe glutamate medium supplemented with uracil at 0.25g/l. The plates were incubated at 32°C for six or seven days until the colonies became visible and were large enough to be conveniently picked.

##### 1.4 Labelling chromosome 2 of CBS2777 with tdTomato and visualization

In order to visualize of the segregation of chromosome 2 in CBS277, we combined the tetO array/tdTomato labelling system of Watanabe and colleagues [1] with the site-specific recombination system described above. Thus we sub-cloned the 10kb tetO array of [1] into the pFA6a-natMX6 REV *attP*$^{\phi C31}$ *attP*$^{\phi C31}$ attB$^{Bxb1}$ integration vector (Figure 2) and then used ϕC31 integrase site-specific recombination to integrate the tetO array adjacent to the centromere in strain Nott373. In order to express the tetR-tdTomato fusion, we replaced the NatMX marker in pNAT ZA31-tetR-Tomato of [1] with HygMX and targeted the Z locus between *zfs1* and SPBP7E8.01 on chromosome 2. The targeted cells were checked for the integrity of the tetO array by Southern blotting. A laboratory strain of *S. pombe* with the tetO array adjacent to the centromere of chromosome 2 at the D107 locus [2]was used as a reference. Both strains expressed *cnp1* N-terminally tagged with mEGFP [3]). For imaging, cells were grown in YEA at 30 ºC and mounted in microfluidic plates (CellAsic. Y04C) with constant YEA media flow at 4 psi. Imaging was performed on a DeltaVision Elite system with the environmental chamber set to 30 ºC. Images were recorded every 10 sec with a CoolSnap HQ2 camera using an Olympus 60x/1.42 Plan Apo N objective, solid state illumination at 461-489 nm and 529-556 nm, a QUAD DAPI/FITC/TRITC/Cy5 multi-band filter for separating excitation and emission, and 525/48 and 597/45 emission filters, as well as 'optical axis integration' across 3.2 µm in z-direction. Chromosome segregation was scored for both the tdTomato-labelled chromosome 2 and all EGFP-Cnp1 labelled chromosomes. In 'symmetric' segregation, red/green fluorescent dots separated symmetrically to the two daughter cells without showing lagging. 'Missegregation' was scored when either both red fluorescent dots segregated to the same daughter, or when green dot intensities were clearly different in the two daughter cells. Single lagging chromatids were scored as 'lagging', one chromosome separating clearly later than the others were scored as 'later'. 'Inconclusive' was used when segregation seemed abnormal but was not easy to fit into one of the other categories.

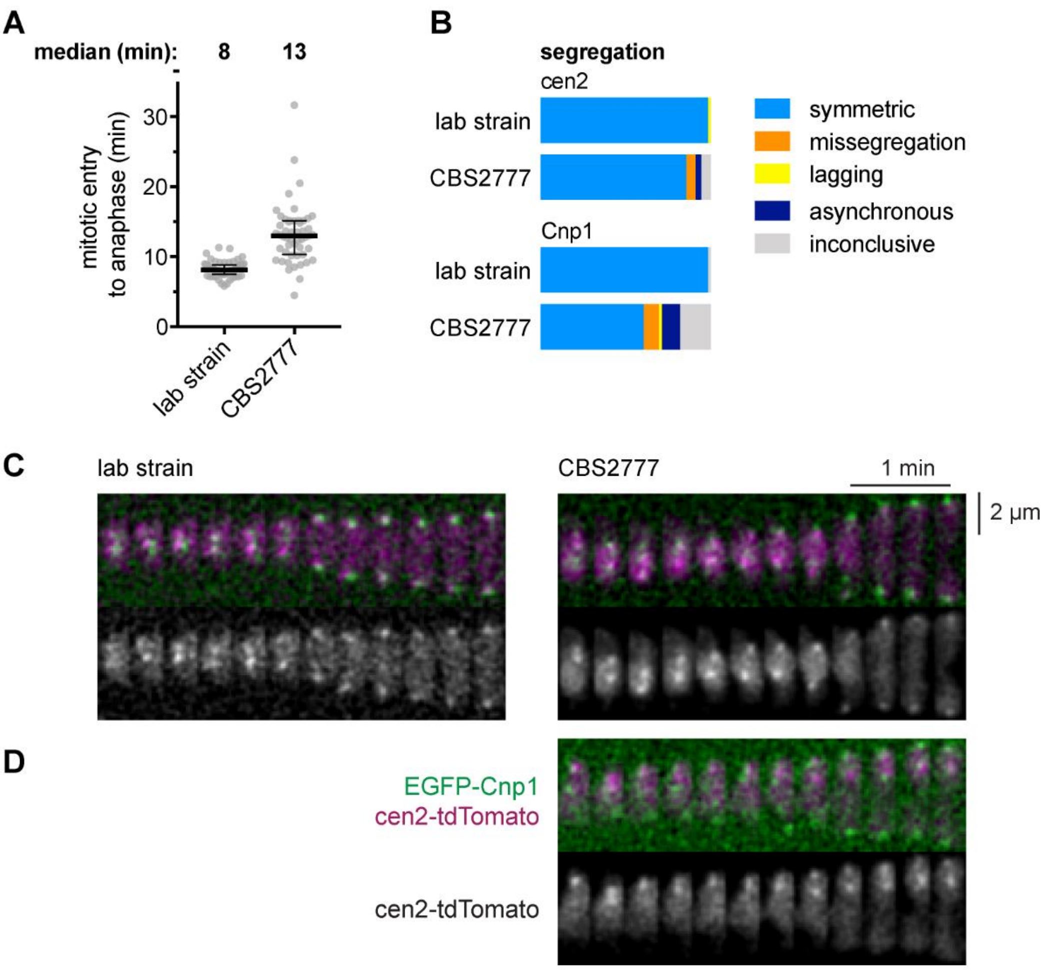


**Supplementary data figure 6. Segregation accuracy of the re-arranged centromere in *S. pombe* CBS 2777**

**A;** Time from entry into anaphase as judged by splitting of CENP-A$^{Cnp1}$ signal for a laboratory strain derivative (SW020) and a CBS 2777 derivative (Nott402) in which chromosomes II and 2 respectively were marked using the *tetO* array/ *tetR* -tdTomato fusion system of Watanabe and colleagues [1] and CENP-A$^{Cnp1}$ was tagged with eGFP. Single cells (grey) with median and quartiles (black).
**B;** The accuracy of chromosome segregation in either a laboratory strain or CBS 2777 derivative in which chromosomes II and 2 respectively were marked as above. The columns illustrate chromosome segregation behaviour (categories as above) of either Cen2 as indicated by the tdTomato signal or the all centromeres signal on the basis of the CENP-A$^{Cnp1}$ ~eGFP signal.
**C;** Imaging accurate chromosome segregation in the laboratory strain and in CBS 2777.
**D;** Mis-segregation of chromosome 2 segregation in CBS 2777.

#### 2 Sequences assayed.

##### 2.1 Central core derived sequences

The sequences assayed in our experiments fell into two classes. One class, of defined lengths was derived from the central core of chromosome II of the laboratory strain of *S. pombe 972* $h^{-}$. In order to help discriminate these candidate sequences from their CBS2777 homologues the candidate sequences were mutated by the introduction of 4bp deletions every kilobase in the 9kb sequence to generate a mutant central core sequence;

```
>mutant centromere
caagcttgtaagcataatatgttcagaactctgtcctttctaatggaaaataatacttatttaagcaatt
ttatcagctacgaacgaccactactacgatgtatgcatgtgaataatatttacatttttttattcctatgt
ctactgtttaaactaagtattgtaaatacttataaaatttattatgatataatgagcttgttctttatt
ttgcaaagcaatatggcttgcatataatacataggctacaattcaatgtacattcaagtattgaaaagct
tttcctgtctttccaatttaaaaacactcaacttcggcgacgtgatataagtataggtataataaataaa
gcgtttttnataaccagnttccgcaaanggtaggaaggtttaatcaattcttaaatggttgaaaagtta
taagaaatagtgatccaattaatcatgccatggaataattttataaaccggtaatcgttgcaaagtgct
taccgntttacttttagggcgaaacaataatacaattaggtagtaccagatcgtttatgaaactgctttt
agataggtactttaaaaccacatgaggtttcagtgaacaacgttttgttatttttttaagtaatgaactta
aactttctttgtttactggttcttatcttactatcataagtattagtaatgtaattttcggtcaaaaaag
gtgcataaaacgacaaaaatgtgttttaaaatttccatttaatttactcatctcatcaattttgtaaagc
taacgaggtatttattttttgcttgtttttatttttaactagtttacgttaaaatttcacttatttagtt
acgatgaatacttgggttaatgtaaaaatggaatgattcgaaacaaaattagttttatcacattcctgtt
ttcgtactttgccttttactatataaataaaaaaaaaaaaaaaaaaaaggaaattgtaacttgaaattt
atgtgattaatttaagcattagcgccttcaataatttatgagtaagatggtaaacaggggtttgattcga
ttataaagggtatggctatcataacttttgttaatccaggcactttttttattgtattctgatttaagagc
tttgatatgatgcatttgaacattttcggtagatttacaaatacaaaacattcttttaattttgatttgg
ttctaattgaccaatatacttcttctcatgaaacattagcgttcatattaaattttatcaaaaatcatca
gttcgaaatcattctacctgtatcacttcaaacaaatgccaaagtttttaacttaacaaaacaaaacgaa
atttttttttgttcttacgtttttcaattactgggggacaaattacgtactaactgttagtttctatg
gtgatttcgttttaattagttactttttttatttattacattcaaaaacacagcttttgctgaacaagact
tgcgtatgaacgcgctaatacaaatataaaaggggttaattttcttcttaattatttacacagaagcgag
cactgtttacatctaacgataatcagaaaattgtgttggcaaaccctagaatgactttaatttttagta
atatgttcttttattgtttagcaaacgtatacttacaaaaggtaattttttactagattttttaaaataga
aatgatctttttgcttaaaagtttaatggtaatattgagatggcccatgactataagatgctattaagta
acgtttatctgaaaatatcattatcattgttcgtatcgcaatttgagatttactaaaccttttttaaaa
tgaaaacgcgaactccgataatcattagtacgaaaatcaaaaaaaaaagttgaagggtacgtttaattctt
ggttttaaaattcgactttagcatcaaagcaaaaacctgaagatttgtatcttattcatttttctttaa
gcggtttagctggttaaagaaaaatgttttccaataatatttattcacatgtactgctttgtgatgcgta
agccgttactcgttaatatgaagaagcattggttagtcctggttgatttctatgtttccttttttggcttc
ggccaattctactttttttaagtattccgtgtgttatctttatggactattattttaaaatctgcatatta
ggtttaagtatgaattagttagggatacgattttctaaatttacatactaataacggaatagatcaaaca
agctccttcaattttagaaaaaaaaaaataacgcattaattgtaaatgttttaggaaattattggtcaa
atctatgaatttgctggttaaatagtatatcaggggggttttatgattttggcgaagtgtcttatccttca
tgttaaatgtattaatttttctttttttcacaaatatcctgtcgattaaaaaattaaaagtaacacatatgt
agtggaatttatctaaaacctcagaattttttaccgagttgtaaacttgacgaatgacatgaaatggtaaa
agtgagcaagctcataaaataaatccacgcttgtcgcaaaacataaagcaaatttgaaaaaaagaagggt
aaatatattaagaatttatttaaaagcaaatcatttaacttacaattgcctaatgtgataactatatcaa
aaatgcccctagcaaagtaggcaacttggttttattgattaacaaaattatgtataagagattaaattat
cctgtttagaaacttactatttctattacacttgctagatagctaagtattaatttatacaaagccataa
tgttgctaacgaaaccttattattgcacattttgacttgaaagggagtctgtaaaccagctataatttaa
ataagactatagcaaatattttagtttataagaaaaatgaatctattaacaacacccgtcatttatgaaa
attaacaacataaatcaaatatgctatatgttaaaattagtactagcgttaaggtaattcgcataagtga
actaagagtcaaagttattttgaacatagttaataaaacaacgatctaatataaattaattaacgttagc
aaggagttttgttccgttcaggtttactgctaacttcaaatcaagaatcaggccattcattaaaaggagc
aatacagaattctaggaaagttacggtttaccagcatattttgaggtacatgaaattttcatcaaacata
ttgtttatctggataggcactaccactaattttaacatgacaatgcgctctaatccatcaatttatcgat
aaatgattgctttcaccactgatttttttatttatttctttttttactatgatagtgccgactaggataa
acgtgtaaattatatcgatgcatttttagaacaagtattactaaatatgcatatcattcaagtaaaaagt
```

gtttgaaaattttaattgcgttagcatattctatttgtcctagtaattgcatctattgtactctctcatc
acgtaaaaaattatttgactacagatagattgctgctgagtaacttattcacacgacgtgcatatccatc
cttcgttgaagaatgcaaaaactcaaattagttaaaatttttaccgaatgtccccccattaaattgaattt
acttaattcagctttctatttccaacgaaaaaaaatattatttccaaaatccaaattaatttattaaata
acatcaatcaaagacgaaaaaagccttattcttcgagaaacgaaacaataaattacctcataattcagtt
ttctattcctaagaccataattcgaataatctttttattaaaatttttaaatttcttcttagtgatttt
acaacacaaaccgactgttagagtagtaggagaagggattgctgtagaaactgctaggtgtgcaaaattt
aaaagataaacttactttatagactatatgtttataactaaatactgactagactttaagttctaccta
ttactggaaaagttatttcctttaacgtaccttttgacagatcctattactggaaaagttatttccttta
acgtaccttttgacctttaagaaacactaaataaaattgaacgtaaataataagtaacgaattctctgaa
ccttttcgataattcatgtgcttgaataaactaatatgtgcaaaaaatcaataccttctttaactcgac
tttataaaatttcccttaagataataattattagcaatgtttaatttgaaagcaaaaattctggtattat
acttgttaagcgctaactcgtttaagtgaaatacggacaaaaaaaaagtaatgaatgcgcagaaaatatc
aaaatacgggtaaattgatttgtttgttaattgccattctttggcgtattgagataattagcaattgcct
tttttaatgcgttcatacggatgactttccacgccatgtcatcaattctttttttttatgtagatacac
ataactcgcatattaactacattagaaattttaaaattaacatatcttgtttgtaatttacaaaccataa
agttttatgataattgttgtgattatcaactttactaatttgactcctaatcagttagaattattttaat
aaatttactaaaacattaaacaaacaacggcacactgtttttttggtcacagcttctaagcatgcaaatga
aattactccaaggaatttgctggcttaataattttatttaataaataaagattatagtagaaaagaatga
aaaagtatttagttgacgatagtatttgttacaaattaaaaactcaaaaccattcttgcagtaagtcaat
cgtgattgacatttctaaaatatcacttactgaacccatgcaagatttagcgcagagggaaacttttata
tctggtagctgagaaacctagtaatgatggtaataaataaatatcaacattcagtttaacatagctataa
acccctaactgtaaccgtagtaaaagctcaaatggctgtatagaagaagttgaaaatcgtttaaagaaac
aatgttgtagtttgaaggatcagttggtaaaacaaatctattatagaaattatattaatttcagagagcc
tcatattttacgcaaattatgaaggtaaatattgcaatcagaccatttgcgaaacagtaaatttcttttt
tctgtttggttttgcttgtgatttggttcatggtttttattaatattttcaagatagtttctaaagatca
tccagaacaatttctattccttcattatttcgttgaaatgaccagatcaaaataaaattctcgaaataat
atttttgcaatttttatgttttattttatatgcaaataaaatgtttacatggaaatcccatatatttaa
aactgactaaagcgtatttaagacgtaatctaggacccctacgtttttttgtttcttgcagttcgaaaaat
cgtgcacatttgtgaaaaggtttagctctacctatcgcaaataaatgctgcatcaaacgcaattgctgct
atgtgggtctgtattgcctctcccttgccagtaatgtgtatttcatcttgtttatttgttttcttatcga
aattaacttaagctttgtccaaatgctaattattaaccactttgggtgtatacgaaacacatattttcaa
aacaccgcaacgaaaaacaaaaaattgtcccttaatatttgttgattaaagtatcgtgaaaggcttttc
ggaaataatctaaaagaatctttaacacctgttaatacttttacaaactaatcgcgagaaaatcattaag
agtttagaataaagtttcctagcaatattttgctatataaatgattttctggttaatggttatagcagaa
aacacagacatttttgtacgtatgtacattaactatgtgaataaacatcttagtacgggttcatcagttc
aataaatattcaactgtattaacaacatgtgcgtccgtgaacatataatgaaagtcaataattatcatga
agctatattttcagttaaaagtttacccagtagatagctctgaagaaatttatttgatgaacagtctaaa
attcaatgcgaaatcctttttaattaatagtgactaatttagtccaagctttcgcgtgtttgcttactttt
attttccctactaaaataaaatctaaaaatttttattcctggaaatcagtaaaactcatgatttcaaaga
aaacagccaaaggtctcagctgctgctagtttgttcgtacgaaaaactgttttcaatattttgactataa
ctagaccactcagattgaatttttttgaaaatttttgtacnttcaaattgcgtcaactgctttaattgtctg
aaatacttgatttgctttcttgagtattcatctagttttatcattaaagctaggtatttccaagggctcc
taaaactaatgattagcgatgtaatatcttaaatataccatttaagttgatgaatgtatagtatgtttta
ctaggatacacggtagaacacatatacttggatgttatgcatacacatcgtagtaattggtccattgttt
ggattgaatgtttagtatgagaactgtatggtggtataccaaaaaatttttacgtttttgaaagtaaaga
tatgattaatatctgcaatgttttcttttagattatggaatctgttacaacagcttaaatggtagtttaa
ttaacaaggtatttactttttattgtttttttgcagctcctccaaaaaattaactagcctagatcacaat
tatcagtcaatctgaattaatagacgtaagaataacttcttaaaaatacagaaaacaaaaaaatgacttc
ataacgtttgtataaaccagatttaattaacggccaataagtttagttacctgttagcaaaaaataattt
taatactgccactcgtcacctttagtaaccctgtaaccgtaattcagcctgtccatcgcaaaaggtaacc
cccatcagtctgtaaatagtaatcttagtagtgcttcaaatattaaaaacggattcgcattctctcaaga
tgtcgaatatgcagaaaaaactttcattattaattattaaacgtaaatctatttacaaatctcaacattt
aaaagttttactaaatataatttaaccataaaataaggttgatgttgaattattctcatgtacaacccaa
aaaaaaggcaatcctacatcattggaatttttaaataaattctttggattcgaaggtctttactcgttt
ttaaaaaaattgatctgaattggcttattagatgacaatatcaaatatgtaccgactcagttgacgttac
ctttatcaaatttatcgcactttaaaacttactaaagcagaagaaaatatgctgaactatttgtggtgga
cattgataatgtaaattacacacttaacgaaattatctagttacaaatatataagctactagtcaataaa
aagctgaaggctcaaatcattacgtaaaatatttttgatcgggaatacaaatactaacttcttgagaaag
ctgcataaaaactagtgtcaatataaaaagcactagtttttcctaatattgactgttgtttacaaatttg
tttactgagcattcagcaaattgcaaaagcctggttaaaaaggaagacgactattctaaatggtatttgt

```
aaaatgcttgctttacttaattaactaataacatagaaaaaactacttttaattttgatcatataactac
agatagaaaaggcagtcaggcaaccagttaacaaccttttagaagtaatgcatacttaagttatgaaaaa
aaaaaattcaagtccacttccaatcccataaaaatgaaataaagcaaacagcagtaaccttgtaaagcac
taaactcattactagaaacagtgaggtgcagggaataaatgtacataatacaaattaagccgcatataag
gacgcagcgagcaaaatagccttcctaatcaaaataactaaaaataaagataaaagaagaaaattcgaat
agtgtaaaattacacacagaaatacttttatgcgagactttcttaggatatgaattatattaactaaata
gcaactgactaatatgtattttgttttgttaagttaaaaactttggcatttgtttgaagtgatacaagta
gaatgatttcgaactgatgattttgataaaatttaatatgaacgctaatgtttccatgagaagaagtat
attggtcaattagaaccaaatcaaaattaaaagaatgttttgtatttgtaaatctaccgaaaatgttcaa
atgcatcatatcaaagctcttaaatcagaatacaataaaaaagtgcctggattaacaaaagttatgatag
ccgtttatacctttataatcaaatcaaacccctgtttaccatcttactcataaattattgaaggcgcta
atgcttaaattaatcacataaatttcaagttacaatttcctttttttttttttttttttttatttatata
gtaaaaggcaaagtacgaaaacaggaatgtgataaaactaattttgtttcgaatcattccattttttacat
taacccaagtattcatcgtaactaaataagtgaaattttaacgtaaactagttaaaaataaaaacaagca
aaaaataaatacctcgttagctttacaaaattgatgagatgagtaaattaaatggaaattttaaaacaca
tttttgtcgttttatgcaccttttttgaccgaaaattacattactaatacttatgatagtaagataagaa
ccagtaaacaaagaaagtttaagttcattacttaaaaaataacaaaacgttgttcactgaaacctcatgt
ggttttaaagtacctatctaaaagcagtttcataaacgatctggtactacctaattgtattattgtttcg
ccctaaaagtaaacggtaagcactttgcaacgattaccggtttataaaaattattccatggcatgattaa
ttggatcactatttcttataacttttcaacatttaaaattgattaaacttcctacatttgcggaactggt
tataaaaaacgctttatttattatacctatacttatatcacgtcgccgaagttgagtgtttttaaattgg
aaagacaggaaaagcttttcaatacttgaatgtacattgaattgtagcctatgtattatatgcaagccat
attgctttgcaaaataaagaacaagctcattatatcataataaaattttataagtatttacaatacttag
tttaaacagtagacataggaataaaaaatgtaaatattattcacatgcatacatcgtagtagtggtcgtt
cgtagctgataaaattgcttaaataagtattattttccattagaaaggacagagttctgaacatattatg
cttacaagcttg
```

Thus it was possible to discriminate between ChIP-seq reads mapping to the native centromeres of chromosomes 2 and 4 of the CBS2777 derived strain and those mapping to the candidate sequences. Subsections of the mutant centromere were amplified with the primers indicated in the table below and sub-cloned into the pFA6a-natMX6 REV *attP*$^{\phi C31}$ *attP* $^{\phi C31}$ *attB*$^{Bxb1}$ vector (supplementary data figure 2). The 0.89kb sequence that was assayed as concatamer was amplified with primers (1240 and 1241) that included BamHI and BglII sites and was concatamerized by sequential BamHI and BglII digestion and ligation. The results of assaying the activities of these sequences are given in figure 3 of the main text.

| **Sequence; numbering with respect to lab strain chromosome II** | **Working name** | **Defined by primers** |
|---|---|---|
| 9.5kb: intact central core | Cen 8 | 247 |
| 6.75kb; 1,620,855-1,627,609 | Cen 5 | 784/785 |
| 4.17 kb: 1,620,825-1,625,025 | Cen 1 | 784/787 |
| 3.58 kb: 1,624,025-1,627,609 | Cen 2 | 785/786 |
| 2.0 kb: 1,624,025-1,626,025 | Cen 3 | 786/789 |
| 1.0 kb: 1,624,025-1,625,025 | Cen 4 | 786/787 |
| 5.48 kb: 1,622,123-1,627,609 | Cen 11 | 785/1412 |
| 4.38 kb: 1,623,232-1,627,609 | Cen 12 | 785/1413 |
| 0.89 kb: 1,624,556-1,625,443 | concatamerized | 1240/1241 |

**Supplementary data table 1**: sequences used in centromere manipulation and assay illustrated in Figures 2 and 3 of the main text

#### 2.2 Candidate sequences

We assayed five different candidate sequences, two of which were derivatives of the 0.89kb sequence that was concatamerized as described above. The first of these sequences is the GC rich sequence which was a concatamer of the following sequence synthesized as a gBlock by IDT. The original sequence from IDT included primer binding sites and was amplified and concatamerized by sequential ligation and BamHI, BglII digestion. This sequence preserves the integrity of A or T tracts greater than three residues in length.

```
> GC Rich
GATCCTTTCGGCGCGCCGCGTATTTTACGCAAATTATGAAGGCAAATATCGTAATCGGCCCATTTGCGAA
ACGGCAAATTTCTTTTTTCCGTTTGGTTTTGCCGGTGATTTGGTTCGTGGTTTTTATTAATATTTTCAAG
GCGGTTTCTAAAGGTCACCCAGAACAATTTCTATTCCTCCATTATTTCGTTGAAATGGCCGGCTCAAAAT
AAAATTCTCGAAATAATATTTTTGCAATTTTTTATGTTTTATTTTATACGCAAATAAAATGTTTACGCGG
AAATCCCATATATTTAAAACGGGCTAAAGCGTATTTAAGCCGTAATCTAGGGCCCCTACGTTTTTTGTTT
CTTGCAGAAAGTCCGAAAAATCGTGCGCATTTGTGAAAAGGTTTGGCGCGGCCTATCGCAAATAAATGCC
GCGTCAAACGCAATTGCTGCTATGTGGGTCTGTATTGCCGCTCCCTTGCCGGTAATGCGTATTTCGTCTT
GTTTATTTGTTTTCTTGTCGAAATTAACTTAAGCTTTGCCCAAATGCTAATTATTAACCGCTTTGGGCGT
ATACGAAACGCATATTTTCAAAACCCCGCAACGAAAAACAAAAAATTGCCCCTTAATATTTGTTGATTAA
AGTATCGTGAAAGGCTTTTTCGGAAACAATCTAAAAGAATCTTTAACGCCGGTTAATGCTTTGCAAACTA
ATCGCGGGAAAATCATTAAGAGTTTAGAATAAAGTTTCCCGGCAATATTTTGCTATATAAACGATTTTCT
GGTTAATGGTTATGGCAGAAAACACAGACATTTTTGGCCGTGTGCGCATTAACTATGCGAATAAACATCT
TAGTACGGGTTCGCCGGTTCAATAAATATTCGGCTGTATTAACAACATGCGCGA
```

The second sequence that we assayed was a concatamer of

```
>In this sequence we have mutated long tracts of A or T by substituting
alternate As with C and alternate Ts with G
GATCCTGTCAGAGAGCCTCATATGTTACGCACATTATGAAGGTACATATTGTAATCAGACCATGTGCGACA
CAGTACATGTCTGTTGTCTGTGTGGTGTTGCTTGTGATGTGGTTCATGGTGTGTATTAATATGTTCAAGAT
AGTGTCTACAGATCATCCAGAACAATGTCTATTCCTTCATTATGTCGTTGACATGACCAGATCACAATACA
ATTCTCGACATAATATGTGTGCAATGTTGTATGTGTTATGTTATATGCACATACAATGTGTACATGGACAT
CACATATATGTACAACTGACTACAGCGTATGTAAGACGTAATCTAGGACACCTACGTGTTGTGTGTCTTGC
AGACAGTTCGACACATCGTGCACATGTGTGACAAGGTGTAGCTCTACCTATCGCACATACATGCTGCATCA
CACGCAATTGCTGCTATGTGTGTCTGTATTGCCTCTCACTTGCCAGTAATGTGTATGTCATCTTGTGTATG
TGTGTTCTTATCGACATTAACTTAAGCTGTGTCCACATGCTAATTATTAACCACTGTGTGTGTATACGACA
GATGTATGTTCACAGCACCGCAACGACAGATACAGCATTGTCACTTAATATGTGTTGATTACAGTATCGTG
ACAGGCTGTGTCAGACATAATCTACAAGAATCTGTAACACCTGTTAATACTGTACACACTAATCGCGAGAC
AATCATTAAGAGTGTAGAATACAGTGTCCTAGCAATATGTTGCTATATACATGATGTTCTGATTAATAGTT
ATAGCAGACAACACAGACATGTGTGTACGTATGTACATTAACTATGTGAATACACATCTTAGTACGTGTTC
ATCAGTTCAATACATATTCAACTGTATTAACAACATGTGCGA
```

In addition, we assayed a concatamer of two sequences in which the CDEII sequences of various *Saccharomyces cerevisiae* centromeres were embedded in segments of the *wee1* gene. The final concatamer is termed “artificial centromere” below and in the main text. The artificial centromere was sequenced by Minion in order to generate the sequences illustrated in figure 5 of the manuscript. The sequences of the units used to generate the concatamer are shown below. The regions of CDEII have been underlined and separated from the *wee1* sequences by forward slashes. In the wee1-CDEII_1 sequence the CDEII and *wee1* sequences were bounded by 17 bp and 19 bp sequences designed to facilitate synthesis and in the wee1-CDEII_2 analogous sequences were 17bp and 10bp in length.

```
>wee1-CDEII_1
GATCCCGTTCTCTAGAG/ATAAATAATTTTAAAAATA/CACGGTTTAGAAATGTTACT/TATTTTTAATAG
TTTTTAAATATTTTA/GGAAGTGGAGAGTTTAGT/GAATATTATTAAAAAGTTTATTA/GCTTACGGCGGT
CCCAACGC/AATAATAATTTAAATTACTATTTTT/CCATGAATCTAAATCGTGCT/TTTTAATAACACTAT
TGTATTTG/GCTCCTCCAACTCCATC/AAAAATTTAAAATACTTTTTTATT/GAGCCTGTATGATGCTAAC
AA/ATTTAATTATTATTAAGTAAAAAA/TTCTACTTCATCTACCTCTTC/TATTAATTAATTTTTTTTCTT
AAA/GCAAAAGCCAAATACCTCTT/TTTGGCAGATTA
```

```
>wee1-CDEII_2
GATCCCGTTGGCTCTAG/TTTATTTTTAATTTTTTTTAA/TCACTAGCTTATTCGGTCCTCG/TTTTTTCT
TTAAATTAAACAAAAA/CAAACAGACCACTTCCTCCC/TATATTTTTAAAAAAAGTAAAAAA/TTCCTTCT
CTCATGCCGCCCT/TTTAAAAAAATAAAATTTAAAATAT/CTTCACCCTCTTTCTCCTCC/AAAATTTTAAA
AAAATTAATTTTCTC/TTTTACACATTCACAACCAGA/ATTTATTATATTTTTTTAATTAC/ATACAGGCA
CAACCTGTACCT/TTTATATTTTTAATTAAATTTTTA/TCATCATCTACTTCACATTTG/TATAAATTATT
ATAATATTGATATTT/TTTTTGATAGACCCAATCTG/AATTAAAAACAAATTATTAATGGTT/GTATCACG
CTCTTCCTCTC/GGATTGGCGA
```

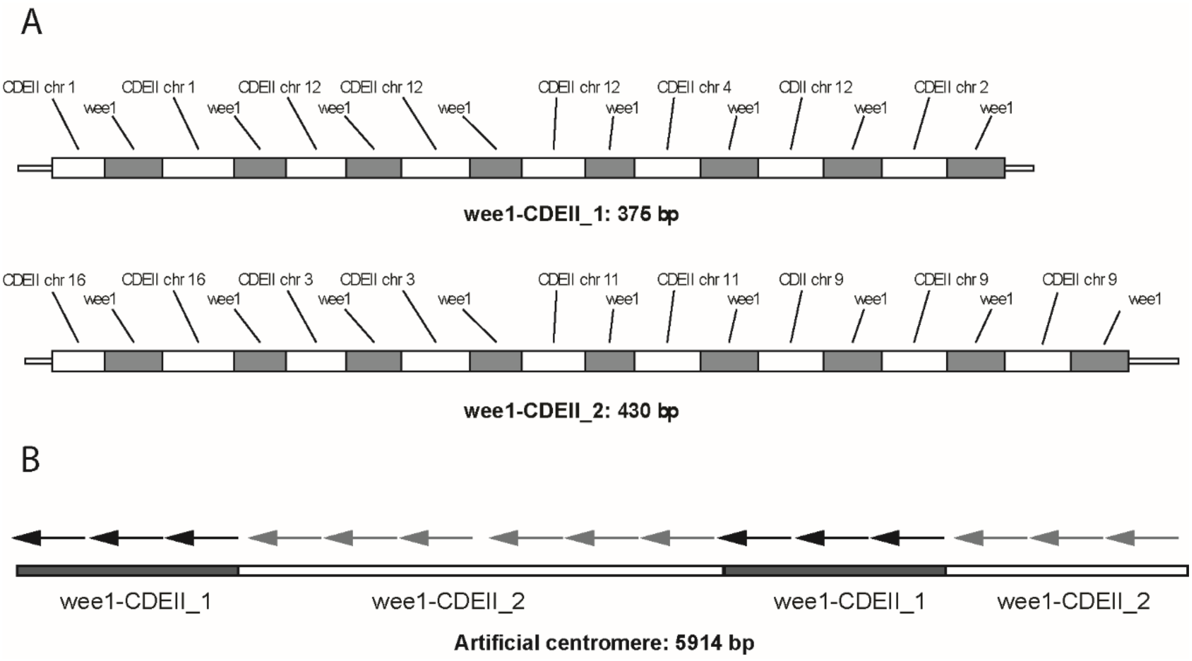


**Supplementary data figure 7; A;** arrangement of CDEII and *wee1* sequences in wee1-CDEII_1 and wee1-CDEII_2. **B;** arrangement of wee1-CDEII_1 and wee1-CDEII_2 sequences in artificial centromere.

In addition we assayed the functionality of the human Y alphoid sequence (GenBank: AF522078.1) and 5.1kb of *Clostridium acetobutylicum* strain LJ4 chromosome (GenBank residues: CP030018.1) residues 3687849-3692959 and 9.0kb of neo-centromeric DNA from CBS2777 chromosome 2 from residues 3862676 to 3871659

>neo-centromere sequence assayed in experiments illustrated in figure 5 of the main text

```
ctgacgacggactttgtgaaggaatcgaatcaaaagatggtttaatcattggtgtacaatggcatcctgaa
gcaatcatcgataaacaaccgcatagcttaaaattatttcaatattttataaaccgctcaaagtggcatat
gaaacaatccgatatatttctaatgttctaccccacgagtcaagttttaccgaaattcaataatctctg
ttccaattgctccctaagtacttgcttgctctttttttttgctgaagtgaattttcggattgttgagaatt
tcagcgttgtcattatgttgtatttaattcaaatggaaagtcctcataattttgcttttttatttatttat
tgcatagtactttctaagtttatatttaattgtgtttatgaaatagagatttcgtcacactatctatatca
gcgaaggattgcaaccgaagagattttagtgtaaaaaatttcgtaaccaccatggcaattctaatctattt
cgttaatcaaattatccaatttattgccaaggttggattatggacatgaaaaaactgtaactcctcaatat
caacactgacaagatactttgtattttaaactatatatttatttatagggggtgcttaataaaattaccgag
aaaatggttattcaagaagcgtgcatgtaatttgtaaagaaaataaattgtggacagaattttaaattaaa
taagcgataagccatcttagacttctcataatcaggcatcttaaatgtatttgggaagcaaaaacttaaat
tcataattttctaaatacattgatctatttaacttaaggagtgaataatgatgatacgtataactcaaagt
gccagtttcctttgcaatgtcattactgtcaaaggaaaaaatgagcaaactttgcttaagtcataaagtat
atataatgcttaaaattatctggccattcaaattttgacttgttctcctttgaagatagatttatgtcgat
aatctttgatacatacttgaatatctgatcatggacataaacagatggcaacgaaagagctcttaattatt
tttttatttttgggagaatagaatatatttgctacggttatccaattggttcatatattggtataaaact
caacatatttaaagtgctataaagataaaaaaattgtatatagcgagcaatatataaatttcatacgatct
aaaatcacgttttaatttctaataccaaaaaaacgataacctttctatatcctaaagttaatagtcaatgt
tttagcaaatagatgttcaattttatatacttgtgaaaatggaaaatcaaattttttttatttaagtaag
aagtgcttataatgtctttgatatcttatcacgatcgaagcttgttttgcatatattcagtccgatttata
aaaaaaattttagttagtagtgcgaattgaagttcttttaataagaactctgctacaaacacacttcagag
aagtttgcatgattcttactatgaccaatctcacacttcagtattttaaagaaacataatccggcggcatc
ttcttaccaaatatattttaaatctgttaatcttggaactaatatgtagctaatttggggtaaatatagct
attatatgataacttgcaataaattatgaacaaaaggaaattgatattttcattaatgtagaaaaaataat
tttcaaaatgatttaatgtctccaataaaaactcgtttaactatataattttctaaatgttgacgatata
ttcatttttaaaaatataacatgtaaccaacagttaatttatacattctaactgcaaaagctataatattc
aacctatcaagtttaaacactaaaaaatctaagatatcttcgaaacataaatataaagctagaagttaatc
acaattatcactccattcttgttgaacaaaataggcaaatgaataaggagaactgtcgtaaccatcatcac
gaacccagtattgtgtaacgtttgaagaagtatataaagtgtcatttgccatattgttcaaagcctttagc
acgatactttgaaggaaattgaggaagacaacatcaattgtttctggcgaggaggtagcattaaacccagc
ttgggcttgaagaatgttaggagtagcatacgaaggaacagaagtatgtactgtatcatctccatcttcac
caaaatcatcacttgtaacataaccgggattaagggcagccaaattttcatttatattgcgtaaattggat
cccccgtttaaggtcgagtataacttcttagcgtatttataaggaacatctgccaaatacatatagtaatt
agtaagaaaagggtttaattggaacatttcgtcgggtccaaattttccttgatacatggaacctcgtatac
cacctccattgctaatgataattcggggcacagttgaacggtttcgattggtaattatttctggcaaaaca
cgatcagtcataagcttaaaaatactgtcatcagattgaggagagacttcagtcatataatagtttttagg
aacgcatccaaagacatatgagaggttaagatcgtcgcgatactgtttaatgattttagataattcaatgc
cttctggagtatcgaaggaatcttctgattgttgagtatggaaacggaagttctggcgattgaaatcaatg
taacggcgcgtgtaatacagaggtgtatttggaatagggagattgggatacgattggcgagtttcattggt
gacgggacgaccaacatattgacgggttgcattggatgctgctaggccatcaattgaaagccagcctacag
tttcacaatatctaccaccctcaagagatacagaagactcgtcgtaaacagcaaagtcacgaatatgagag
tgaccgccaaagatttgaatgggagtatcaggatgcacttttctaattgaggcatgtagtgacttccattc
gtcccaatcgcgaaccggaatatgtccaagtaataagaaaaggtcaacatctgttctatttatttgttgtt
gataccatctggagttgacggctgtctcaacaggtgtaacaacagtgttgttggcgttaccggtgaaatcg
tagaggaagcctacagcaagaactcgaacgccatgtttcgtagtaaagtaggcagattcggctgcaaattg
ctccaactcatttgatgagttgaagatttgaacgttagaagcgagataagttccattccagtgaggtacaa
aatattcgtgggtattattagaaacggatgcttggtacaattcgtgattaccaatcgtaagaatatcatag
ggtaggtacgtaaaaatgttgtttgtatatattccttctggatcagaagcatcagaaagtccgtttccgtc
atgtaaatcaccggtatcaactaataataaatctacgtctttaaaatcagctaattccttcatccgcaaaa
caaaagacttaaattcgccaaagtcggctttataacgagaatctctaaggtgaccaccaagccaaccgtga
gtgtccgtagtatgaatgaaattcatttgaccccattctaaaggtttaatgatttggttccaatcatacat
tgtagttaaatcggtagacgagtaagcaattacagaaccaatacaactgaacaatagcacaaggttgacc
```

aaaaatgtatcgaggctgtcttcatgttgatggagtcgggataaagagcttaacaatctatttcgctgttt
accgagggtaagattctttaatttatacaaaaaaaagatcggcatttcgaaattcaatgttttcaaatttc
cctcagtcaattcattaacatcgtacaacctttgaaaagtgtcagtgtcaaaaaaaggtttacaaaacttt
gtcggagatttcagtcccattttgctaatgacctaatgtcaaaatttagtaatgaattgaagttgcgaagc
aactttgagtttgctgcaaatgtatcttcttacattatattaatttaaacaatcaaaattcataataatta
tagacagtgttatttcttcgttcccttactcaaacatttatttaagtaaaaaaactctttagcaatgggaa
tgactcggagttcgatgggaaaaccgtaacatcagtctcaaaatactgttaatggaacatttgtgttttgc
attatgtcatttacgtccttcggtatataataacgctatgttcggttgacaaaaagttgtcctaaaagttt
atttttaagacatgcactttctgcttacgatttaaaaaaaaaaaaaaacattgtcttgctaagctataatt
gaatattttgtatgagctaagaattattgttagagctgtgagaaccagttctagtacttcaatgagcttct
actatgattgttgtccatattaaacattgtatcggattcaatctctccgggaaacaccaagccgagcggac
catgaatgagagcatacagactttcgattctgttcattacgaaaaatattaggtagtacatagccgacaaa
tcacttaccggtaggatatgccaatatcgacaagcgtatgggatggtgcaaaaacggtaatatttcaaagt
tggcttcaataatagtcgacaaaagctctaagtttaaaagacgaagtttaataacatttgtgatggtttgc
ctttttacatcaagagaaatttcattatgcatataaaacaaaagcatatttgaattaatcaaaataacca
attattgcttagaaccaaagaaaccccgatttacgcaattaacttaaataaaattattttgcaaggaatat
tcgactttcgaatggaagtattttaacttcgacgaacattgaaatgacattgcaagcatttccgttcaaat
tgttcaatattttataaaagtcagaagttaagattagaactaatatattgactactttttttgactactt
caacttaccggagctagactactccgacattgcaaaacgtgagattggtttatcaaatatttaccacttta
atatcagttattttcaaaatatcttttactttatttgcaattgctaaaagctagttttcgaaatgaaattc
atcattggaagttagagcttaatatctagtccgtcactgagttcttcatgaagtctgtcgtacattttgag
agataaagaatatgaaatttgtcatttatgttgcttgtttgcccaaattagcattttattgacggtaaagg
ataatgacataataaaaaagagggatgggatagaagcggaagtagtattgctcggcatattaatatgattt
ttcaagagttaagacggtttatggtatataacttcagcgagctaagcacattaaattcaccaggaattaag
cagaactacaagaaggaattctttaagaaattttggataaggtgacttttaccaatatattttctgaaatc
acatattcaaatttgtctaaaaaaagaataacagcaaaaaaagcatacagtgtactgtccaccaatattac
taaagaaagtatgagcagaaacaatcatgaattgatccaacatcgtctagacattgcgtaaaagcaataaa
atataaaattgaacttaaagaacctttcatattgacaataaagtatcattaatgtagacggaaaagatac
aatactaaaacttttgaaagggttcaaaaaaattgagtaaatacttctatttaaccattaacccattcaa
ttcgattttaagctggagtcacagataaaataagtaagtgattacatattctgtttgaacacactcatgtc
gcatgttctgtaaaaaaggttgaaatttaaaatgtagtattttgagtgatcattggacattaatgctaact
tactctttaacttaattcctaagagggagtatactttttctttaagatatccgtaaaatatagctttctag
ttctctatttctcgacattattaaggttcacattcttaaaagttacaaacatctttaaaattgtaaaact
ttttaagatactatagagttcattaacttttattaaataagtatatagtaaataaccgaactctgaaaact
gcagctttcgttttccggaaaatagcaacaaaagaatttgattacaaaaatgagaattcataaaaaaaaca
aatgaagctatgaagggtgaataaattctacagccttgtcgcaatgacatgttggtctactctatctacga
tcgtactcgtagacacagtagtaatatttcaaacaagtaaaaattaaacgatcagcaacttaccaacatgt
acctttcattttgactgtgtttaatagtgtatttttacattactattaagaatcgagtatatatcgatga
gttcaaaattctgtaatatcaactcaaaagacttggaccaaagatagcaacaaaattacaaataaaccaac
gctattaacaagtctccaaattccagaataagctctgtttatatccggctggaaaaatctccttaactgca
aatttttcctttgtgcaaataaaatcacagaaaattaaaaaaattagtaacgattattagacctttctttt
ttcacgagggtaatgctgataataaagttcagttattaagagatcagtttcctttgaataatcctatcttt
cttttagtgaacaacaaactaagatattttctgatatattttaataataatgaattcagactatatacata
aatgggcaatgaatccgaatagagatggttaaatgcaaaacaatgatgaaagcgagaacagaaagacatat
tttccataaaatcaataacatactttgtaattttttgttagctttaagattgattgttgtctctaattgaaa
gggtttccccgttgcgagtaggtagagtatgaaaggcaatctcatctttaaatggttcaatctttaagtcc
aaagtcaatattactactttatagagagcccaatggctgaaatacaaaagaccggttatagttgcacccaa
ggctgctataatcattcccaaggctgcacccaggactgctatcatcataccggtctcccgtaaagcaagca
ccacaaactctgcagagactattttgaattattagcaaaaaaaaaggcatcactgtaaaggaaacatacaaa
ttaaactaacaaacaataaaacaaacaaacaaatgtaaatagcaataggccccccaaatttcactgtaaaac
cgattttcatttataaaactttcatttatccaaattaaaagaaaatctttgacttcaaagttcttgtctat
tgcgaaaataattaaaatacaccaaataataagcaagggccaacgaagtttactgcgggtagaaatcctac
catcaggataacaactatcatgaaatttagaaatttcagtaaaaaggttatttaaaaagacctttattta
tctgtcttaatgtttataatatcttcttgaggcttgatggtgttggtaaagagattttgtttgtttacctt
ttcaggcttacacaattcaagtccccgctccaggtcaacattacgagtatctcttaccataaataactcat
tgtaaccaggaggttcaacttgtttttttgggctttctggatttgacattggaataaaaccacagcactgc
aatttgacagctttagcaccactagtaatgtttgacaatataaaattaaacagtgtggtttatggaagata
ataagaaaatataataaaatattttaagccataaacgaaagaattaaaaaaagatcaaaattggtgaatgt

```
tttacgtatccttaaatcagataccaaactgcgtagctttctttattataccgaaaacattcacactgtct
acaaaaattactaagaatactataggctgcgatgagttttcttagtgattttttgttttttttaaggttta
cacattgaatgaaaaaaaaaaaacagtatatgtagcatcaataggacttaagcatttcgacagatttaat
atttattacgaaaatgttcatcgaaagtagtagaagatcttccaaatatcttcagattcctttgattcgta
caaggactaataagaccatcttcaaagaaaaaataagaattattttaaagatgtcaaacatccgcataca
attgttttgtatcgggatggtattgtttatttattagtttcagaactttattatctaaagttgagtaacaa
tactataaatcgtttttgcaattgaatatgccgtgaccatttcaaatgctgttagcgtgaaaagcatggga
acaaacagaccgcagacaattagatataaattcaagatcagagaaaaggaagaagtgattgatgattgtt
tacccaattggttcaatgactttattcaaattaggaaattaggccattttaatattaacttctgacgaaaa
gcgggagaatcaaccagtgttgtaaatttgtcttaagtcgatgatatttgcaatgttcacgcaatttagat
gaatttttcgcttcgctataagaatgatacatgggcaatcaaccctaacccatagtgcgtagccaatcatt
ttagttcatgctgtcgcaaaacagaaagggaattgcaaacccttagcagtagtaagtaaacccttgttat
aaataatgcttgtacatgaatccggttatgttgaaaggactatttttattcatttgatttgggataatga
ttgtacaacataaaacagcaaaaatagaagaagatcatggcttattccaaccaatcttgcgtccctcagat
atcagcaaaacaactgatacgaaatttattcagtcttccccgtatattgaaaaagagcattggctggactt
gggaaccttaagtgtcggccattacttttatcactagctcttcagacttttgttcccaaagattctgtcc
gttatgcccatcttccttatgctcaagcctttgatatt
```

#### 3 Chromatin immuno-precipitation (ChIP)

##### 3.1 ChIP

ChIP was carried out as previously [4] using Abcam AB290 anti-GFP antibody and Dynabeads Protein_A conjugate (Invitrogen / Thermo Fisher 10001D) for the precipitation of the GFP tagged Cenp-A and anti-c-Myc-Agarose Affinity gel (Sigma A7470) for the precipitation of the Myc tagged Cenp-C

##### 3.2 Computational analysis of ChiP-sequencing data

The raw reads were firstly processed using the Sickle software for quality trimming and the Scythe software for PCR adaptors trimming. The trimmed reads were 75 basepairs on average. The BWA-mem software was used for all the alignments with standard parameters, to the genome *Escherichia coli* strain K12, sub-strain MG1655 (GenBank accession number U00096) to remove possible contaminants.

The genomes of the engineered strains were built on Vector NTI® software using the previously sequenced genome of the start strain CBS2777 [4] and the sequence of the plasmids used for the engineering. The reads unmapped to the *E. coli* genome were aligned, using BWA-mem with standard parameters, to the correspondent *S. pombe* genome. This resulted in primary mapped reads. Then the reads mapped in correct orientation and within insert size were selected using samtools "flags". Unmapped reads were removed from the bam files and the total number of reads mapping to each centromere were then determined using the following chromosome coordinates: chromosome I: 2926051-2939353, chromosome II: 125168-133320, chromosome III: 1092658-1106291, chromosome IV: 1619693-162784.

When necessary, the number of reads covering the neo-centromere on chromosome II was determined using the following coordinates: chromosome II neo-centromere: 3855830-3867330.

Coverage vectors were created for using the R package Bioconductor, then coverage plots were created in R. The plots were smoothed using the default running medians scatter plot smoothing in R. For coverage plots across entire chromosomes data points were plotted every 1000 base pairs, and to plot the centromeric regions data points were plotted every 100 base pairs.

##### 3.3 Results of ChIP-seq analysis discussed but not illustrated in the main text

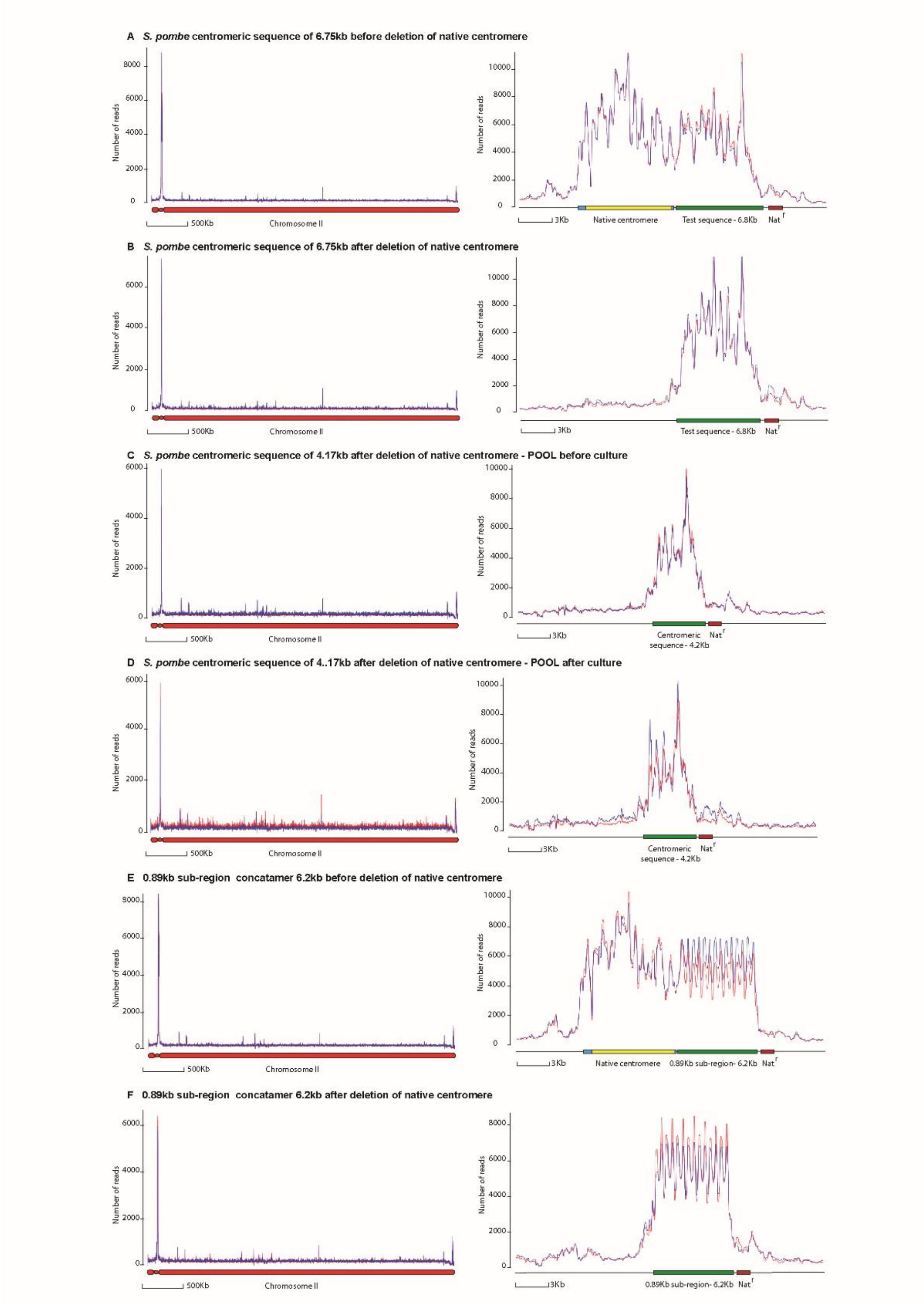


**Supplementary data figure 8;** ChIP-seq analysis of centromere replacement on 6.75kb sequence and a 6.2kb tandem repeat of a 0.89bp centromeric sub-sequence. A: 6.75kb sequence adjacent to native centromere prior to deletion of the native centromere. B: 6.75kb sequence after deletion of the native centromere. C: pool of 4.17kb sequence after deletion of the native centromere. D: pool of 4.17kb sequence after deletion of the native centromere and long term culture E: 6.2kb tandem repeat of 089kb sub-region adjacent to native centromere prior to deletion of the native centromere. F: 6.2kb tandem repeat of 0.89kb sub-region adjacent to native centromere after deletion of the native centromere.

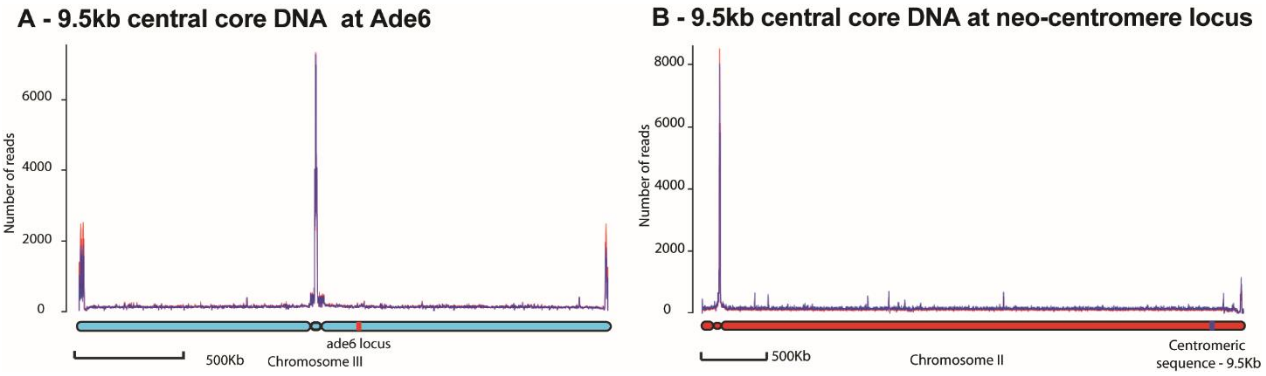


**Supplementary data figure 9.** Absence of ectopic binding of CENP-A$^{Cnp1}$ to centromeric sequences when placed at non-centromeric loci; **A;** 9.5kb of central core DNA placed at Ade6 locus on chromosome 3, **B;** 9.5kb of central core placed at neo-centromere locus on chromosome 2.

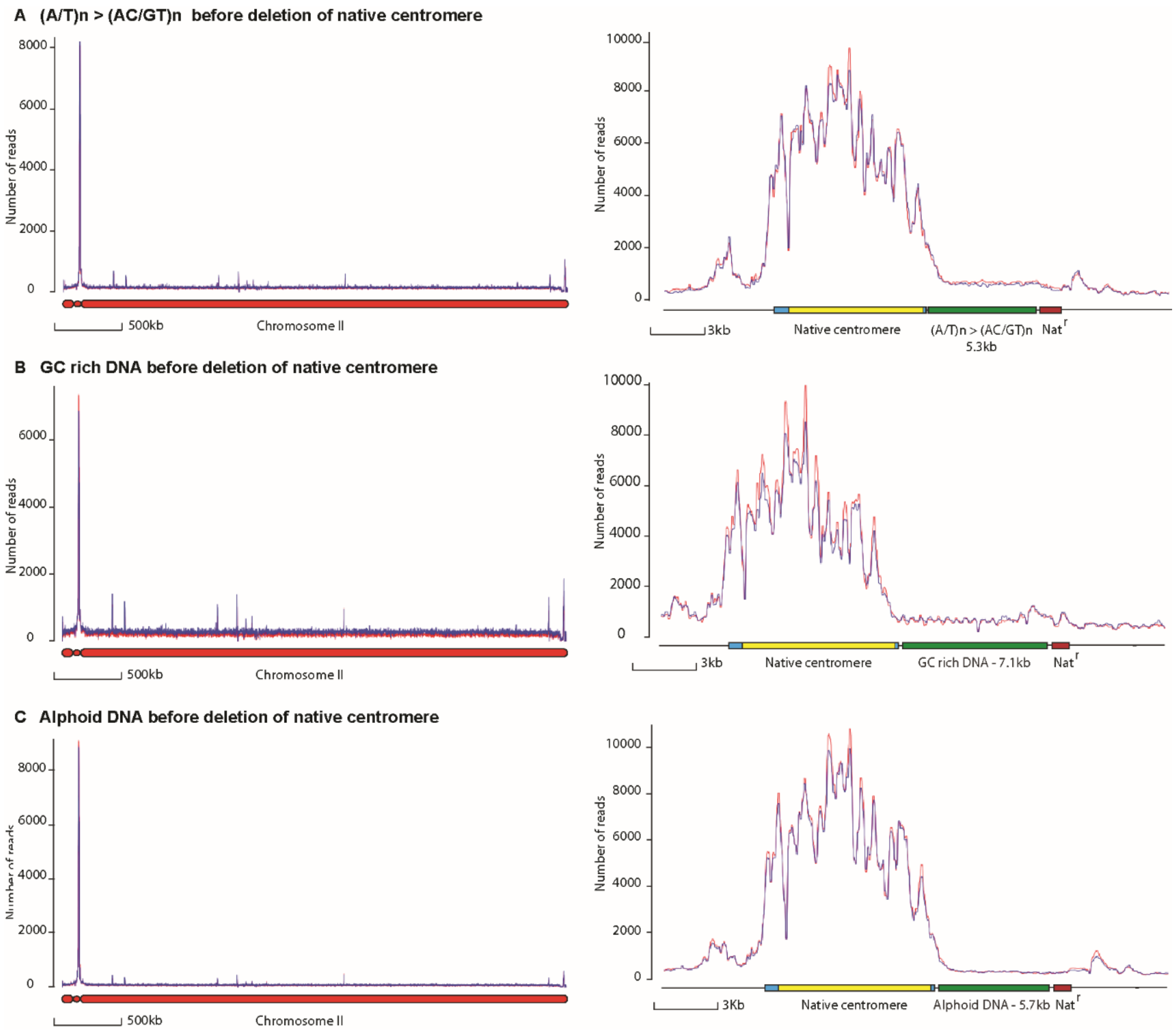


**Supplementary data figure 10.** Absence of ectopic binding of CENP-A$^{Cnp1}$ to GC rich derivatives of the 0.89kb sub-section of the central core or to human Y alphoid DNA centromeric sequences when placed adjacent to the native centromere of chromosome 2 of CBS2777.
